# Spatially tuneable multi-omics sequencing using light-driven combinatorial barcoding of molecules in tissues

**DOI:** 10.1101/2024.05.20.595040

**Authors:** Giorgia Battistoni, Sito Torres-Garcia, Chee Ying Sia, Silvia Corriero, Carla Boquetale, Elena Williams, Martina Alini, Nicole Hemmer, IMAXT consortium, Shankar Balasubramanian, Benjamin Czech Nicholson, Gregory J. Hannon, Dario Bressan

## Abstract

Mapping the molecular identities and functions of cells within their spatial context is key to understanding the complex interplay within and between tissue neighbourhoods. A wide range of methods have recently enabled spatial profiling of cellular anatomical contexts, some offering single-cell resolution. These use different barcoding schemes to encode either the location or the identity of target molecules. However, all these technologies face a trade-off between spatial resolution, depth of profiling, and scalability. Here, we present <u>B</u>arcoding by <u>A</u>ctivated <u>L</u>inkage of Indexes (BALI), a method that uses light to write combinatorial spatial molecular barcodes directly onto target molecules in situ, enabling multi-omic profiling by next generation sequencing. A unique feature of BALI is that the user can define the number, size, and shape, and resolution of the spatial locations to be interrogated, with the potential to profile millions of distinct regions with subcellular precision. As a proof of concept, we used BALI to capture the transcriptome, chromatin accessibility, or both simultaneously, from distinct areas of the mouse brain in single tissue sections, demonstrating strong concordance with publicly available datasets. BALI therefore combines high spatial resolution, high throughput, histological compatibility, and workflow accessibility to enable powerful spatial multi-omic profiling.

## Introduction

The physical location of a cell and its interaction with its surrounding microenvironment profoundly influence its gene expression, regulatory networks, cell identity, and function. Prime examples are the concerted activation of homeobox genes in response to morphogen gradients in embryonic development^1,2^ or the role of Notch signalling between neighbouring cells in controlling embryonic and adult neurogenesis^3^ as well as synaptic plasticity^4^. Yet, our ability to discover new and more fine-grained examples of spatiotemporal control of biological functions has been limited by the resolution and scope of technologies for deeply profiling the molecular make-up of cells while maintaining spatial information.

The emergence of methods for spatial omics offers the ability to interrogate the identity and function of cells in the context of their local tissue environment. Among many other findings, these technologies contributed to unveiling spatially defined axes of cell differentiation during gut development^5^, provided insights into patterns of interactions between maternal and foetal tissues during placental development in early pregnancy^6^, and metrics to refine the stratification of cancer patients for improved treatment outcomes^7, 8, 9^. What was once a technological void is now crowded with solutions that strive to find a balance between the depth at which molecular information is profiled and the resolution of the associated spatial coordinates.

Key to all existing methods is the use of a barcode to either identify target molecules when spatial coordinates are detected by microscopy (e.g., MERFISH^10^, SeqFISH^11^,/SeqFISH+^12^, StarMAP^13^, in situ sequencing^14,15^, FISSEQ^16^, ExSeq^17^), or to encode their spatial coordinates when next generation sequencing is the detection method (e.g., Spatial Transcriptomics^18^, Slide-Seq^19^/Slide-SeqV2^20^, Slide-tags^21^, HDST^22^, Stereo-Seq^23^, Pixel-Seq^24^, DBiT-seq^25,26,27^, Light-Seq^28^). The method to assign and decode barcodes defines the characteristics of the technology in question. Optically decoded barcodes offer superior spatial resolution since they preserve tissue histology, yet this comes at the cost of lower measurement depths, as fewer distinct molecules can be profiled simultaneously. On the other hand, nucleic acid barcodes allow an unbiased and broad coverage of gene expression since the target molecules are read through sequencing, although usually with a coarser spatial resolution due to various technical limitations in the way the barcode is produced and linked to biomolecules. Several techniques adopt a grid-like distribution of spatial barcodes^18–27^ that is independent of the tissue’s histological features, posing challenges in accurately capturing the tissue’s structural characteristics. While in some cases the distribution of the barcoded features in the grid can offer sub-cellular resolution (i.e., HDST^22^, Pixel-Seq^24^, Stereo-Seq^23^), this comes at the expense of throughput or of the ability to profile large areas of up to many centimetres squared. Furthermore, methods using spatial DNA barcodes do not allow the pre-selection of regions of interest within the tissue, resulting in sequencing reads being wasted on profiling areas not relevant for the question at hand and further reducing information depth. There are currently few technologies that combine genome- or transcriptome-wide coverage with the fine spatial resolution necessary to profile the molecular phenotype of single cells, and existing approaches do not scale easily to a throughput sufficient to dissect how their interactions within tissue neighbourhoods respond to perturbations during development or pathogenesis in large sample cohorts.

The ability to integrate different types of measurements (e.g., mRNA, chromatin, and protein) allows for a more comprehensive characterization of cellular identity and function. Multi-omic integration can be achieved either by projecting data collected in parallel from comparable samples onto a reference model^29^, or by measurements from the same physical sample^30,31^. A common strategy for accommodating multi-omic profiling on a single physical sample is to transform the biochemical diversity of different target biomolecules into a unified chemistry for barcode assignment^32,33,34,35,36^. For instance, Share-Seq^31^ and DBiT-seq^26,27^ both employ a nucleic acid handle that can be attached to many types of biomolecules to enable subsequent serial ligations to generate a barcode that encodes the cell identity or spatial location, respectively. While these techniques hold promise for capturing spatial multi-omic information, they often require custom-made, complex devices or other specialized equipment, limiting their broad accessibility. Moreover, in many cases these workflows do not easily integrate into standard protocols for sample preparation and processing, putting them out of reach for many labs.

In response to the limitations of existing approaches, we developed Barcoding by Activated Linkage of Indexes (BALI), a spatial multi-omic sequencing method employing user-defined optical delivery of spatially referenced molecular barcodes directly to target molecules within intact tissues to enable their identification, measurement, and localization by next generation sequencing. To achieve this, a patterned illumination by UV light directs the spatial linkage of barcode subunits by DNA ligase through serial cycles. Each cycle adds short oligonucleotides representing the individual digits of combinatorial barcodes that are appended onto the target molecules. The assignment of spatial barcodes is spatially tuneable from millimetres down to micron scale (∼5 μm), and it is mostly limited by the technical specifications of the light patterning method. Depending on the complexity in the combinatorial barcode, the number of areas interrogated is similarly scalable, ranging from one up to millions. Our current efficiency of each cycle of ligation is ∼86%, which corresponds to an overall ∼48% efficiency in encoding a thousand spatial locations. We characterized the efficiency of the ligation dependent barcode encoding both on solid supports and tissue sections and assessed the resolution of light directed ligation. As a proof of concept, we used BALI to profile the gene expression of two different areas of the embryonic mouse brain and benchmarked it against RNA sequencing after physical dissection by laser capture microdissection (LCM)^37^. Additionally, we profiled chromatin accessibility of different areas of the adult mouse brain using a 4-digit barcode, finding strong agreement with publicly available datasets. Lastly, we combined gene expression profiling and ATAC-seq to obtain a multi-omic characterization of two areas of the adult mouse hippocampus. We demonstrated that BALI barcodes can be written with subcellular resolution and high efficiency, theoretically allowing the rapid profiling of thousands of areas. Thus, BALI has the potential to open up new opportunities in spatial biology, offering a histology-aware approach to multi-omic profiling of tissues with tuneable resolution, throughput, and layering of genomic information.

## Results

### Barcoding by Activated Linkage of Indexes

At the core of BALI is the use of light to drive sequential ligation of individual oligonucleotide modules to encode combinatorial spatial barcodes directly on intact tissue samples. Initially, tissue sections are placed on conventional histology slides and fixed / permeabilised (Fig. 1a). As a first step toward installing the spatial barcodes, a universal ligation root adapter is coupled to target biomolecules in situ. The ligation root is an oligonucleotide with a short (6 nt) single stranded DNA overhang and a photocaged phosphate at its 5’ end (1-(2-nitro-phenyl)-ethyl based linker)^38^. Importantly, this can be installed on target molecules as part of conventional workflows, including reverse transcription and tagmentation, to profile mRNAs and genomic or epigenomic features in a single experiment. Once prepared, the tissue is imaged and segmented into spatial barcode locations of interest, as defined by the user. These that can range in size from large tissue regions to sub-cellular compartments, and in number from few barcodes to millions. Each location is assigned a specific spatial barcode which is then encoded through cycles of illumination and ligation. Upon illumination, the photo-cleavable linker is released to expose a 5’ phosphate group for ligation to the molecule encoding the subsequent barcode digit with a defined, compatible ligation overhang.

**Figure 1.**
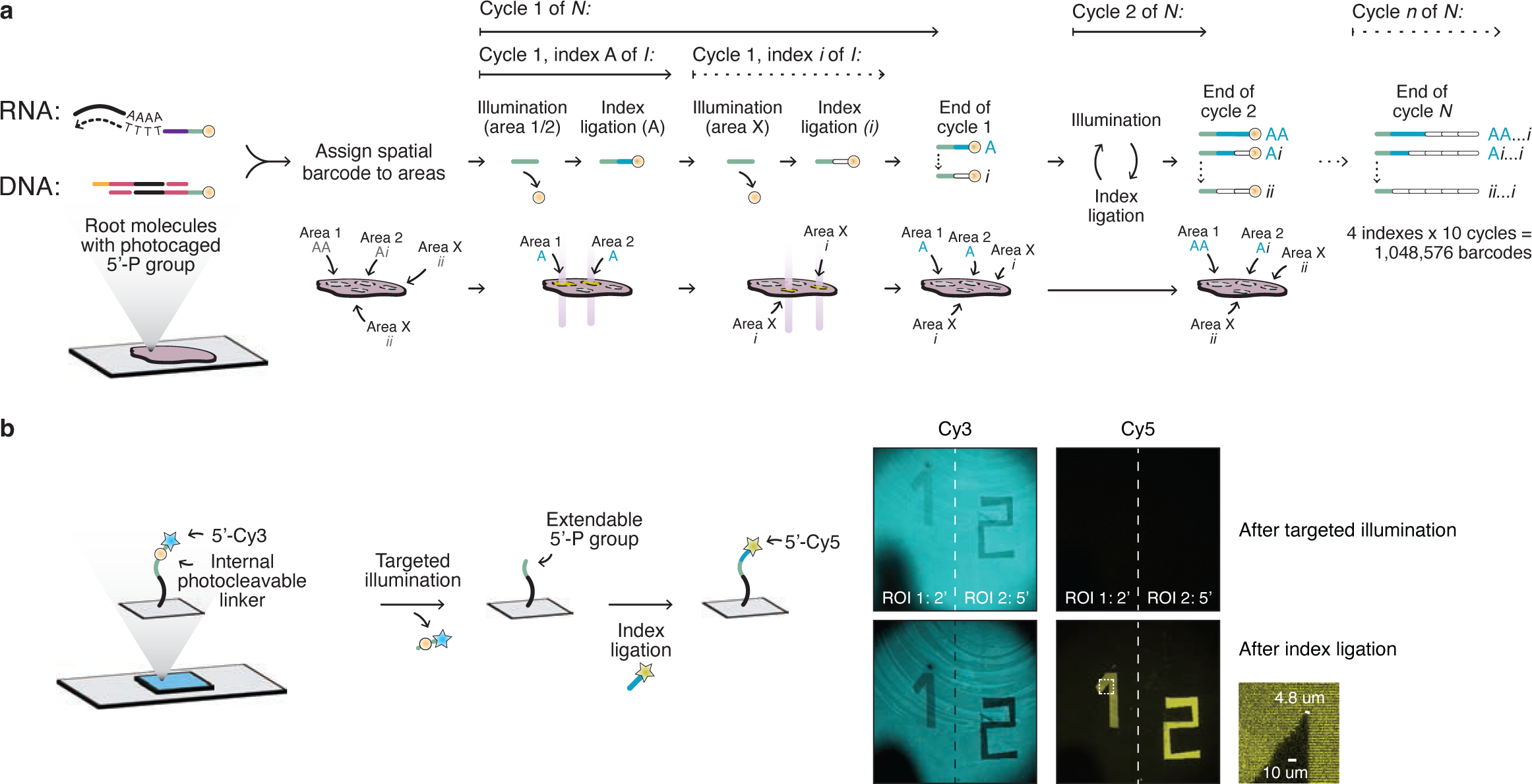
BALI is a technology for spatial-omic profiling that combines high throughput and high spatial resolution. a) *Schematic workflow*. Thin tissue sections are placed on glass slides. After fixation and permeabilization, reverse transcription or Tn5 tagmentation are used to couple each target molecule with a ligation root equipped with a 5’ overhang and a photocleavable cage on the 5’ phosphate. The tissue is imaged, and multiple areas of interest are defined by the user. Each area is assigned a unique multi-digit molecular barcode, composed by a sequence of short DNA “indices”, that will get encoded through cycles of targeted UV illumination and ligation. In the first cycle, all the areas with value ‘A’ in the first digit are simultaneously illuminated to release the cage from the 5’ phosphate, to make it available for ligation with a small dsDNA that encodes the ‘A’ index. The cassette regenerates an orthogonal ligation overhang with a photocleavable cage. This process is repeated until all areas receive their first digit, and then again until all digits are encoded in all areas. The resulting molecules are extracted, amplified, and sequenced. Bioinformatic processing can map each molecule on the genome or transcriptome and in physical space. b*) Quantification of BALI barcoding resolution on a solid surface*. A schematic of the experimental set up and microscopy images of spatially resolved ligation. A slide was covalently functionalised with oligos featuring a 5’ terminal Cy3 fluorophore attached through an internal photo-cleavable linker. Upon UV illumination of a region of interest for 2 minutes (ROI1) or 5 minutes (ROI2), the photo-cage was released producing a 5’ terminal phosphate group, and the oligo thus made available for ligation with a staggered dsDNA cassette featuring a 5’ terminal Cy5. The images show the signal of the anchored oligos (Cy3, on the left) and ligated cassette (Cy5, on the right), post UV illumination and post ligation (top and bottom rows, respectively). An enlarged detail of the internal corner of the “1” shape shows that the method can achieve a spatial resolution of at least 5 μm. Images are in false colours in a colour-blind friendly palette.

By providing different variant DNA molecules for each digit and using light to control in which region each is installed, it is possible to write complex barcodes combinatorially. The length and complexity of the barcode is determined by the desired number of spatial regions that will be distinguished in each sample. The total number of theoretically possible barcodes is defined as *i^N^*, where *i* represents the number of different identities for each digit of the barcode (also referred to as the barcode’s base), and *N* represents the number of digits in the barcode. For example, a 10-digit barcode with 4 different values for each digit is sufficient to identify ∼ 1 million (4^10^) distinct spatial regions. The base or the number of digits can be further increased to implement codes with error detection and error correction.

During each cycle, all the spatial locations assigned to the same value *i* for the digit *n* to be encoded are illuminated to make their physical spatial barcode molecule extendable (Supplementary Video 1). A ligation event appends the index molecule encoding value *i* and regenerates a ligation overhang with a photocleavable cage to the 5’ phosphate, available for writing the following digit. Cycles are repeated to encode all the different values for digit *n* of the spatial barcodes, before moving to following digit and so forth, until the barcode is fully assembled, and all the spatial locations receive a unique DNA identifier with *N* digits. Indexes in the last cycle append a Unique Molecular Identifier (UMI) and an additional primer annealing sequence for PCR amplification. The resulting libraries are prepared and analysed via sequencing by synthesis with standard 300 cycle reagents (Supplementary Fig. 1a): read-1 (100 bp) is used to map the molecule on either the transcriptome (RNA) or genome (chromatin), whereas read-2 (200 bp) decodes the spatial barcode.

Each digit of the barcode is encoded by a staggered dsDNA molecule, here defined as an “index”, with a specific DNA sequence that is different for each digit value (supplementary Fig. 1b). The indices are flanked by two orthogonal single strand ligation overhangs on both 5’ ends, each with a different sequence and length (6 or 7 nt). On the strand that extends the ligation root, the ligation overhang includes a photocaged 5’ phosphate.

As a proof-of-concept, we functionalized a glass slide with a ligation root oligonucleotide carrying a Cy3 fluorophore on its photocaged 5’ phosphate. Illumination of the slide with a focused UV laser (405 nm) for either 2 minutes (Region of Interest 1, ROI 1) or 5 minutes (Region of Interest 2, ROI 2) resulted in cleavage of the photocage and release of Cy3, as evidenced by the reduction of Cy3 signal in illuminated regions. The removal of Cy3 exposed the 5’ phosphate, which we then used as a substrate for ligation with a short Cy5-labelled dsDNA molecule to mimic the addition of the first digit of a barcode. Both uncaging and ligation were strictly confined to the illuminated areas (Fig. 1b), and the relative signal of Cy3 and Cy5 was dependent on the amount of light received (Supplementary Fig. 2). By measuring a sharp corner of the illumination mask (the smallest resolvable feature in this image), we determined the resolution to be ∼5 μm, therefore compatible with sub-cellular measurements. These results indicate that ligation can be controlled with high spatial resolution in situ.

### Optimal design of ligation overhangs

In order to optimise the design of our indices, we investigated the dependency of the activity of T4 DNA ligase on the structure of the DNA ends to be adjoined. First, we optimised the length of the overhangs to balance the requirements for strong annealing during ligation while minimising off-target binding, something often observed with long single stranded DNA molecules. Across different experimental conditions compatible with BALI, we found that the optimal length for the ligation overhangs was between 6 and 7 nucleotides.

Secondly, we assessed the sequence dependency of T4 DNA ligase activity. We had observed significant differences in the efficiency of ligation for different overhang sequences, independently of the overhang length and conditions of ligation. Such dependency had been previously described for shorter sticky ends (4 nt)^39^, and thus we resolved to expand this analysis to overhangs 6- and 7-nucleotides long. To this end, we screened two individual libraries covering all possible 6-mer and 7-mer overhang sequences to identify those that were able to drive the highest ligation rates in solution (Supplementary Fig. 3a). The screen showed an inverse correlation with between the GC content of the ligation overhang and the activity of T4 DNA ligase. The top 10 hits from each library (GC content > 30%) were individually validated both in solution and in situ for their ability to drive efficient ligation. The validation confirmed the overall ranking of our screening results (Supplementary Fig. 3b, 3c). Furthermore, we assessed the fidelity of ligation for the top scoring candidates by ensuring no cross-reactivity between different overhangs. This was particularly relevant to avoid the formation of concatemers from the same indices. The top scoring 6-mer and 7-mer sequences were selected as alternating ligation overhangs.

### Efficiency of barcode encoding by iterative ligation

Combinatorial barcode encoding by serial ligation is key to the high-throughput nature of BALI. As an initial proof-of-concept of this, we tested whether we could write a 4-digit fluorescent code, in which each digit could have two possible identities (Fig 2a, Supplementary video 2). A 5’ photo-caged ligation root was functionalised on a glass slide and subjected to serial cycles of illumination and ligation to extend a DNA barcode. Each alternative index molecule, here defined “A” or “‵B”, carried a different 5’ terminal fluorophore (FAM or Cy3). This allowed us to track the addition (and subsequent uncaging) of index molecules visually. All 16 barcodes could be successfully encoded, as shown in Fig 2a.

**Figure 2.**
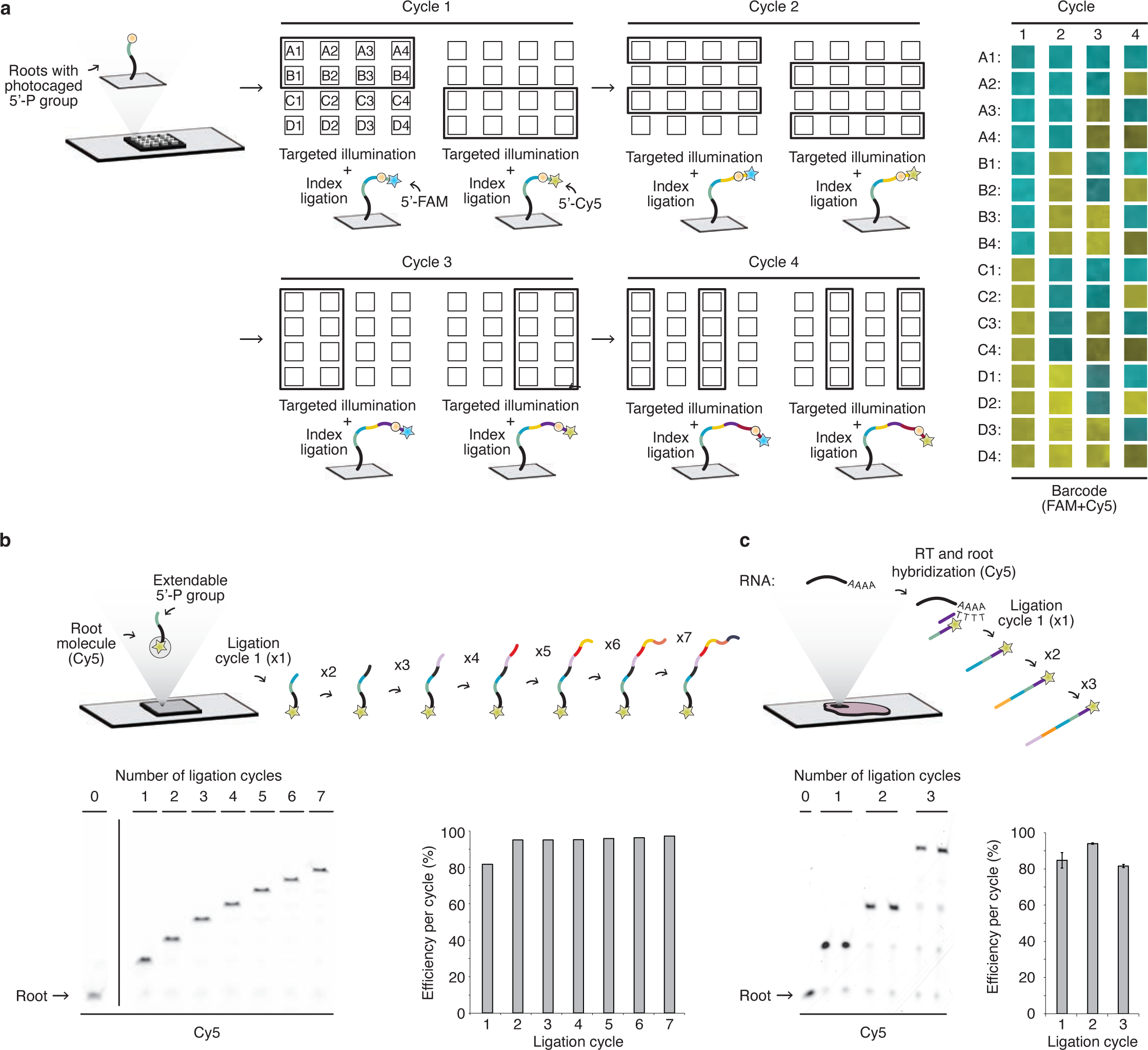
BALI efficiently encodes multi-digit spatial barcodes. a) *Fluorescent barcodes encoded on a solid surface though 4 cycles with 2 different indexes.* A schematic of the protocol (on the left). An oligo with a photocaged 5’ phosphate similar to the one described in Fig. 1 was immobilised on a functionalised glass slide to serve as the ligation root for sequential cycles of UV illumination and ligation with fluorescently labelled dsDNA indexes (FAM or Cy5) producing all possible combinations (4 digits ^ 2 variants = 16). Imaging was performed after each cycle and represented in false colours (panel the right) for all the 16 areas (rows) and cycles (columns). Images are in false colours. b) *Quantification of sequential ligations on Sepharose beads*. A schematic of the protocol (top panel): an oligo featuring a 5’ phosphate and 3’ Cy5 was annealed to a complementary oligo immobilised on Sepharose beads. The fluorescent oligo was extended through multiple (n=7) cycles of ligation with dsDNA indexes. The efficiency of each ligation step was measured by densitometry of the Cy5 signal on a denaturing polyacrylamide gel (bottom panel). c) *Quantification of sequential ligations on thin tissue sections (mouse liver).* A schematic of the protocol (top panel): poly-A mRNA transcripts in thin tissue sections were detected by hybridization with a poly-dT tailed oligonucleotide. This served as molecular docking for the same fluorescent ligation root described in Figure 2b. The root was extended through multiple (n=3) cycles of in situ ligations. The efficiency of each ligation step was measured by densitometry of the Cy5 signal on a polyacrylamide denaturing gel (bottom panel). Error bars represent the Standard Deviation of two replicates.

The fraction of complete barcodes produced in a BALI experiment depends exponentially on the ligation efficiency of each cycle (Supplementary Fig. 4a). Therefore, high ligation efficiency in each individual cycle is critical. To measure the individual and cumulative efficiency of barcode writing in our system, we encoded a 7-digit barcode on oligonucleotide-functionalised Sepharose beads (Fig. 2b). First, the beads were hybridised to a ligation root consisting of a 5’ phosphate and a 3’ terminal Cy5 fluorophore, which was used to identify the molecules on an acrylamide gel. The efficiency of each ligation cycle was measured by densitometry of the bands at each iteration, while the cumulative efficiency was measured as the product of all the previous cycles. After 7 cycles, the cumulative efficiency of ligation was 60.6% with each individual cycle ranging between 81.4% (cycle 1) and >94.8% (cycle 2 through 7). The lower efficiency in cycle 1 was consistent across replicates, which may indicate difficulty in performing ligations in proximity to the bead surface. We also performed a similar experiment to measure the efficiency of barcode ligation in fresh frozen tissue sections from adult mouse liver (Fig. 2c). In this case, the fluorescent ligation root was hybridised to a poly-dT oligonucleotide primed on poly-A tails of cellular mRNAs, and we tested three cycles of ligation. The efficiency of ligation was consistently above 80% for all three ligation cycles in each replicate (ranging from 80.6% to 93.8%), with a measured cumulative efficiency of 64.8%. This efficiency can be extrapolated to a forecasted efficiency of 48%% for a 5-digit barcode or 23.1% for a 10-digit barcode, which would label a thousand or million locations, respectively (Supplementary Fig. 4b). BALI is therefore able to encode multi-digit spatial barcodes on both solid supports and biological tissue samples to label a large number of areas with high efficiency.

### Spatial RNA profiling of mouse embryonic brain regions during neurogenesis

One of the fundamental attributes of BALI is its ability to tailor both the quantity and arrangement of the spatial profiling regions according to experimental requirements. The significance of this adaptability is two-fold: first, it optimises sequencing costs by focusing exclusively on relevant locations and, secondly, it adopts a histology-driven approach to spatial definition, ultimately allowing high-throughput profiling of meaningful locations rather than arbitrary coordinates.

To demonstrate an end-to-end BALI workflow for spatial transcriptomics profiling, we applied the technique to two functionally and anatomically distinct areas of the mouse embryonic brain: the subventricular zone (SVZ) and the developing cortex (Fig. 3a, 3b). In this experiment, the ligation root was introduced during in situ reverse transcription and template switching using a dedicated reverse transcription primer that consisted of a poly-T region at the 3’ end for priming on the poly-A tail of mRNAs, and a photo-caged ligation root overhang at the 5’ end. Targeted UV illumination of the SVZ released the photo-cage from the 5’ phosphate selectively from the cDNA molecules within that region. These molecules were then ligated with an index consisting of a short, staggered dsDNA molecule harbouring a defined sequence. The same process was repeated on the cortical region to ligate a second index, producing a simple 1-digit barcode on each region. After ligation of the spatial barcodes, cDNA was purified from the lysed sections and PCR amplified with dedicated primers to generate libraries compatible with Illumina sequencing.

**Figure 3.**
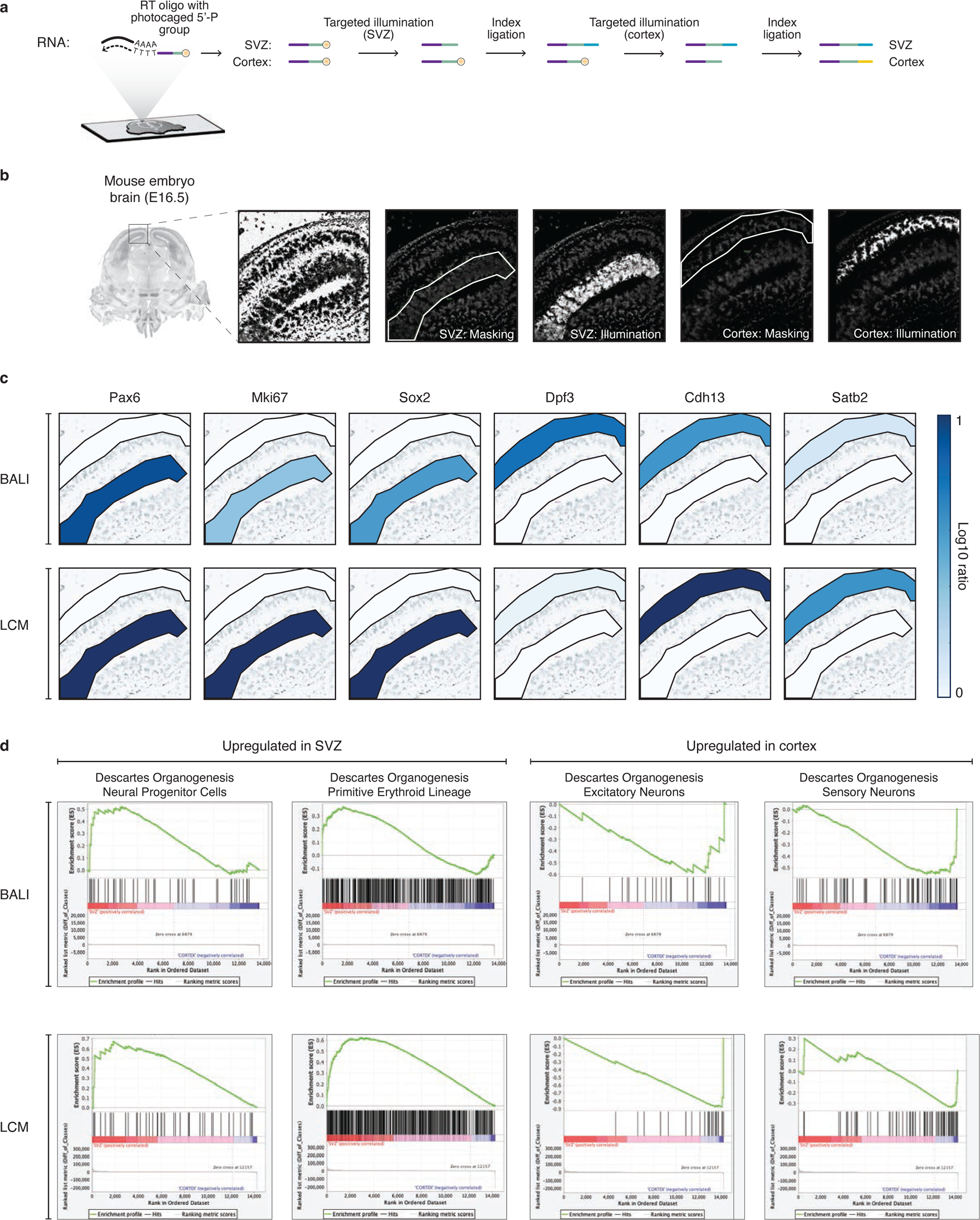
Gene expression profiling of two functional areas of the embryonic mouse brain. a) *Schematic workflow.* Thin tissue sections from fresh frozen embryonic day 16.5 mouse brains were placed on conventional charged microscopy glass slides, formalin-fixed and permeabilised. In situ reverse transcription and template switching were performed with an RT oligo comprising a poly-dT-VN 3’ end region and a 5’ terminal ligation overhang (green) ending with a photo-caged 5’ phosphate (yellow circle). The resulting cDNA was therefore linked to a photo-caged ligation root. The sub-ventricular zone (SVZ) and cortex of both hemispheres were sequentially UV illuminated to release the photo-cage and ligated with index ‘a’ (blue) and ‘b’ (yellow), respectively. b) *Representative microscopy images for the tissue during uncaging.* The coronal full section overview is from the Allen Brain Atlas for E16.5 mouse embryonic brains^56^. The close-up panels are from our samples (one hemisphere only). The first panel shows the nuclear stain from Draq-5 upon excitation with the appropriate 647 nm laser, and it is shown as black on white. The following panels show the illumination masks used (white outline), and the resulting excitation with UV laser of the Draq-5 stain. c) *Gene expression of a curated list of markers.* The expression levels of 3 marker genes known to be upregulated in the SVZ (left) and cortex (right) are shown in a scale of blue (log10 ratio) over the relevant illumination masks. Top and bottom rows represent the expression levels as measured with BALI and LCM (with bulk RNA-seq), respectively. d) *Gene expression of gene panels through Gene Set Enriched Analysis.* Gene-Set Enrichment Analysis (GSEA) outputs for 4 gene sets from the Descartes gene signature collection^41^ in the SVZ (left) or cortex (right) using the expression levels measured with BALI (top row) and LCM (with RNA-seq, bottom row), respectively.

Following sequencing, we were able to detect 5348541 and 1930824 UMIs for the SVZ and cortex, respectively. Given an average cell diameter of 12 um2 for this tissue type, this corresponds to an average of 484.6 UMIs/cell (707.6 UMIs/cell and 261 UMIs/cell in the SVZ and cortex, respectively).Notably, this is within the range achieved by commercial solutions such as 10x Visium on a comparable sample40 when scaled for a similar binned area of (50 um2 diameter) (BALI = 12284.2 and 4542.1 UMIs/cell vs 10x Visium = 6212.8 and 16504.3 UMIs/area in the SVZ and cortex, respectively) (Supplementary Fig. 5).

Next, we investigated the expression of a curated panel of marker genes and confirmed the enrichment of cDNAs associated with neural progenitors in the SVZ (e.g., Pax6, Mki67, Sox2) and differentiating neurons in the cortex (e.g., Dpf3, Cdh13, Satb2) (Fig. 3c). In addition, Gene Set Enrichment Analysis (GSEA) revealed enrichment of biologically relevant signatures from the Descartes gene signature collection41, such as ‘neural progenitor cells’ and ‘primitive erythroid lineage’ in the SVZ, and ‘excitatory neurons’ and ‘sensory neurons’ in the developing cortex (Fig. 3d).

To benchmark the results obtained with BALI, we performed Laser Capture Microdissection (LCM) followed by bulk RNA-seq on the same brain regions from an adjacent slide. The results from LCM closely matched those obtained with BALI, both in terms of individual marker genes and GSEA outcomes (Fig. 3c, 3d). Therefore, we show that BALI accurately captures the transcriptional profiles of specific spatial regions and yields results consistent with traditional methods such as bulk RNA-seq on physically dissected tissue.

### Spatial chromatin accessibility profiling of mouse adult brain

To expand the portfolio of information accessible through BALI, we applied our method to profile chromatin accessibility within the adult mouse brain. This was achieved by combining in situ Tn5 tagmentation with BALI’s iterative cycles of ligation/illumination on two distinct brain regions: the dentate gyrus and the cortex (Fig 4a, 4b). In this experiment, the design of the ligation root was slightly modified. A directly accessible 5’ phosphorylated ligation overhang was appended to one of the two adapter oligonucleotides loaded into Tn5, without any photo-cleavable group present at this step. We introduced this modification to streamline the pre-processing of the tissue, eliminating the need for light-protected conditions during tissue preparation. To thoroughly assess the protocol’s compatibility with multi-digit spatial barcodes, a unique 4-digit barcode was assigned to each region. After tagmentation, the first ligation introduced a first shared barcode and an orthogonal ligation 5’ overhang featuring a photo-caged 5’ end to all fragments across the tissue. Subsequently, two cycles of bulk illumination and ligation appended the second and third digits of the longer spatial barcode. The fourth digit was then ligated upon targeted illumination to either the dentate gyrus or the cortex.

**Figure 4.**
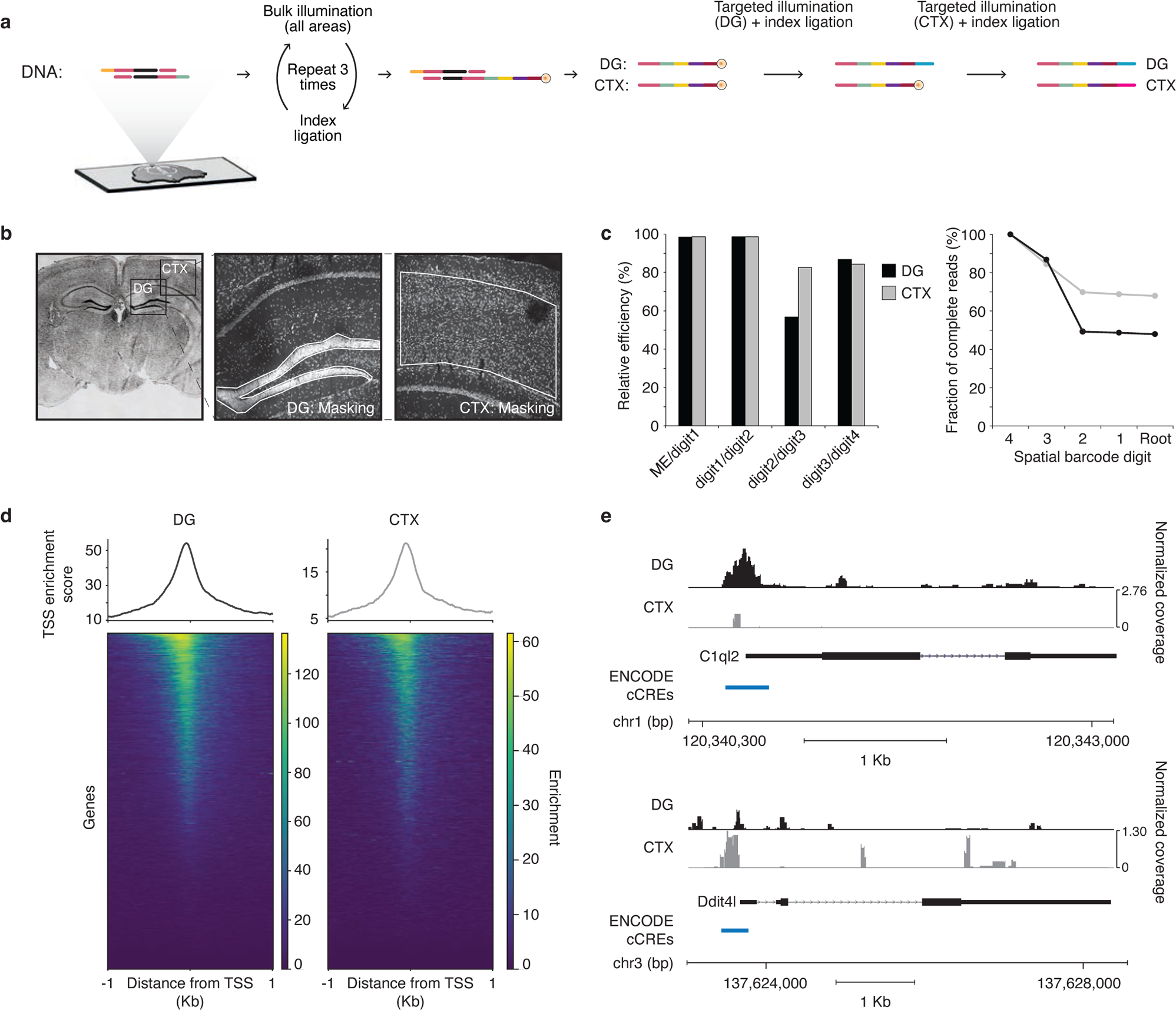
Chromatin accessibility profiling in two areas in the adult mouse brain. a) *Schematic workflow.* Thin tissue sections from fresh frozen adult mouse brains were placed on conventional charged microscopy glass slides, fixed and permeabilised. In situ Tn5 tagmentation with custom adapters targeted the accessible regions of chromatin. The resulting tagmented fragments were flanked on one side by a conventional A-adapter sequence (yellow) ^57^ and to a 5’ ligation overhang (green) on the other. Sequentially, three rounds of ligation and bulk UV illumination extended the root with three different photo-caged indexes (yellow, purple, magenta) to encode the first, second, and third digits of a 4-digit spatial barcodes. The resulting 5’ end carried a 5’ photo-caged phosphate. The Dentate Gyrus (DG) and a region of adjacent cortex (CTX) were sequentially UV illuminated and ligated with area specific indices (blue and orange). b) *Representative microscopy images for the tissue during uncaging.* A coronal full section overview of the tissue section, with annotation of the fields of view for uncaging (overlayed boxes). The close-up panels identify the illumination masks (white outline) used for uncaging. Signal in all panels refers to Draq-5 nuclear stain. c) *Efficiency of ligation for the multi-digit barcode for each area.* Efficiencies (defined as the fraction of full-length barcodes detected) are shown as relative percentages for each individual digit (bar plot, left) and cumulative rates (line plot, right). Black refers to the DG, grey to the CTX. d) *Enrichment at transcriptional start sites (TSS).* The data is shown as cumulative plots (top) and heatmaps for individually ranked TSSs (bottom), for the DG (left) and CTX (right). The graph shown refer to 1 kb upstream and downstream of the TSS. The scales are adjusted for each area. Black refers to the DG, grey to the CTX. e) *Representative coverage plots showing differential chromatin accessibility across two genomic loci.* Accessibility for the DG and CTX are shown separately in black and grey, respectively, for C1ql2 and Ddit4l. The RefSeq annotation is shown below, as well the annotation for gene promoters (blue) from the ENCODE’s cCREs database^46^.

Next, we lysed the samples to extract the tagmented fragments and generated libraries for sequencing via PCR amplification with dedicated primer sequences. The frequency of spatial reads that included the Mosaic End sequence and a complete barcode was 47.9% and 66.9% for the dentate gyrus and cortex, respectively (Fig 4c). Furthermore, we investigated the rate of occurrence of incomplete barcodes, such as those lacking one or more digits due to failures in intermediate ligations. Given the alternate pattern of ligation overhangs across digits, the most abundant incomplete barcodes were 2-digit barcodes, consisting of digits 3-4 or 1-4, which derived from failures of either the first or second ligation, respectively. Notably, failures in the first ligation were 15-30x more frequent than in the second ligation, recapitulating our prior observations on Sepharose beads.

The quality of our ATAC-seq datasets was assessed across a range of different metrics. We obtained a fraction of reads in called peak regions (FRiP score) of 0.48 ± 0.03, a TSS enrichment score of 24.05 ± 5.01 and a percentage of reads mapping in mitochondrial genes of 6.44% ± 1.04%. These values were all within the range of the data standards set by the ENCODE consortium^42^, and were comparable to those reported for DBiT-seq in brain tissue^43^. The enrichment of fragments in proximity to Transcription Start Sites (TSS) (Fig. 4d) showed a skewed distribution immediately preceding the TSS, consistent with open chromatin in promoter regions. Notably, we could detect differential chromatin accessibility in the dentate gyrus and in the cortex. In particular, the promoter region of C1ql2 has been previously shown to be a marker locus for the dentate gyrus in a single-cell ATAC-seq atlas of manually dissected brain regions^44^. Our data shows a similar chromatin accessibility profile that is specific for the dentate gyrus (Fig. 4e). We also investigated the promoter and enhancer regions of genes that are differentially expressed in the cortex and dentate gyrus based on the Allen Brain Atlas IHC dataset^45^. Consistent with the known gene expression levels, we detected increased chromatin accessibility at Prox1 and Lhfpl2 loci in the dentate gyrus (Supplementary Fig 6a, 6b). Similarly, Pamr1 and Ddit4l loci exhibited higher accessibility in the cortex (Fig. 4e and Supplementary Fig. 6c). The peaks of accessibility overlapped with promoters and proximal or distal enhancer regions as defined by the modENCODE consortium^46^, emphasizing the accuracy of the chromatin accessibility profiles we could obtain from this BALI-ATAC workflow. This data underscores the ability of BALI to profile chromatin accessibility across multiple spatial locations in intact tissues with a histology-driven approach.

### Spatial multi-omic characterization of the adult mouse hippocampus

To ask whether BALI could perform spatial multi-omics, we next attempted to profile simultaneously the transcriptional and chromatin landscape across two distinct regions of the adult mouse hippocampus in thin tissue sections. Combining the transcriptome and chromatin profiling workflows explored above on the same tissue section, we first performed an in situ chromatin tagmentation step to install ligation roots on open chromatin, immediately followed by in situ reverse transcription and template switching to incorporate primers encoding the same ligation root in cDNA derived from polyA mRNAs (Fig. 5a).

**Figure 5.**
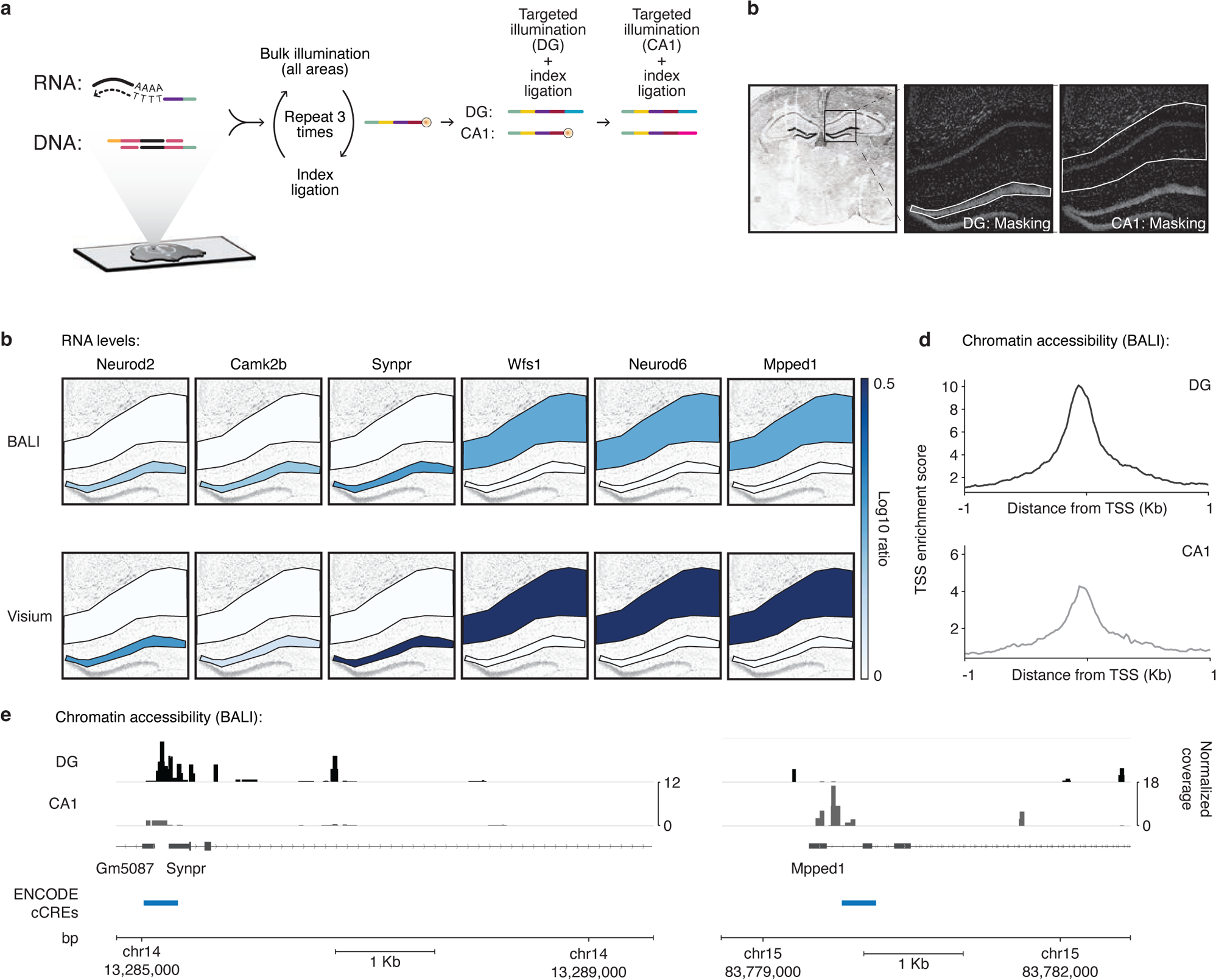
Multi-omic profiling of gene expression and chromatin accessibility in the adult mouse hippocampus. a) *Schematic workflow.* Thin tissue sections from fresh frozen embryonic day E16.5 mouse brins were placed on conventional charged microscopy glass slides, fixed and permeabilised. In situ reverse transcription and template switching followed by Tn5 tagmentation with custom oligos were used to detect and label the tissue mRNAs and accessible chromatin with the same BALI ligation root with a 5’ ligation overhang (green). Sequentially, three rounds of ligation and bulk UV illumination extended the root with three different photo-caged indexes (yellow, purple, magenta) to encode the first, second, and third digits of a 4-digit spatial barcodes. The resulting 5’ end carried a 5’ photo-caged phosphate. The Dentate Gyrus (DG) and the CA1 hippocampal region (CA1) were sequentially UV illuminated and ligated with area specific indices (blue and orange). b) *Representative microscopy images for the tissue during uncaging.* A coronal full section overview of the tissue section, with annotation of the field of view for uncaging (overlayed box). The close-up panels identify the illumination masks (white outline) used for uncaging for the DG and CA1. Signal in all panels refers to Draq-5 nuclear stain. c) *Gene expression of a curated list of markers.* The expression levels of 3 marker genes known to be upregulated in the DG (left) and CA1 (right) are shown in a scale of blue (log10 ratio) over the relevant illumination masks. Top and bottom rows represent the expression levels as measured with BALI and a previously published 10x Visium Dataset^47^, respectively. d) *Enrichment at transcriptional start sites (TSS).* The data is shown as cumulative plots (top) for the DG (left, black) and CA1 (right, grey). The graph shown refer to 1 kb upstream and downstream of the TSS. The scales are adjusted for each area. e) *Representative coverage plots showing differential chromatin accessibility across two genomic loci, Synpr and Mpped1.* Accessibility for the DG and CA1 are shown separately in black and grey, respectively. The RefSeq annotation is shown below, as well the annotation for gene promoters (blue) from the ENCODE’s cCREs database^46^.

Following tagmentation and reverse transcription, we performed iterative cycles of ligation and illumination to encode the appropriate spatial barcodes in either the dentate gyrus or the CA1 region of the hippocampus, with the same encoding scheme as described for our ATAC-seq experiment (Fig. 5a, 5b). After sequencing, we were able to detect overexpression of Neurod2, Camk2b and Synpr in the dentate gyrus and Wfs1, Neurod6 and Mpped1 in the CA1 (Fig. 5c), in line with publicly available data profiled with combined 10x Visium and scATACseq^47^ (Supplementary Fig. 7, 8). As observed in our single modality spatial ATAC-seq, chromatin accessibility at genic loci showed a peak immediately upstream of the TSS across all biological replicates (Fig 5d, 5e). Chromatin accessibility was differentially distributed in the two regions analysed. Among the differentially regulated peaks, we confirmed differential chromatin accessibility for two loci for which we detected matched differential gene expression. First, we identified dentate gyrus-specific peaks in the promoter of Synpr, a gene encoding a synaptic protein known to be exclusively expressed in the flossy fiber of the dentate gyrus and CA2-CA3 of the hippocampus^48^. Conversely, chromatin accessibility at the proximal enhancer of Mpped1 was restricted to the CA1 region (Fig. 5f). These data demonstrate the versatility of BALI for spatial profiling of tissues to collect multi-omics information from a single sample.

## Discussion

Here we have described BALI, a method for multi-omic profiling of intact tissue sections via light-driven combinatorial ligation of DNA barcodes with user-defined resolution and throughput. While some existing spatial sequencing methods similarly utilize light activation to define regions of interest (e.g. Light-Seq^28^ or GeoMX^49^), BALI introduces a combinatorial encoding scheme that exponentially expands the throughput range from tens to millions of independent spatial locations – thereby enabling single cell analysis in situ.

Importantly, BALI can achieve subcellular-scale spatial resolution (∼5 μm), which could be improved by using higher resolution optics than the 10x objective currently used. While other existing spatial omics methods exists in which spatially referenced barcodes are arrayed with high resolution (up to 2 μm for HDST or 10X Visium HD), BALI barcodes can be assigned computationally to areas defined by the user, which can allow for non-uniform shapes and sizes to capture unique and important features of histology and biological knowledge. Combining light activation with combinatorial barcoding, BALI offers a solution to the conflict between resolution and throughput that has thus far dominated the field of spatial profiling. Together, these features enable BALI to offer both high-throughput profiling and histology-driven analysis of intact tissue samples.

BALI’s reliance on serial ligations to generate combinatorial molecular indices makes it inherently constrained by the efficiency of ligation. This is a bipartite problem. Firstly, it requires efficient tagging of the biomolecules of interest with an oligonucleotide that could be the substrate of ligation events. The ability of BALI to capitalize on existing priming and tagmentation workflows to install the ligation root provides robust efficiency at this initial step. Secondly, the efficiency of barcode creation multiplicatively depends on the efficiency of each subsequent ligation step. Interestingly, we showed that the first ligation reaction consistently experiences the lowest efficiency, which could be due to steric hindrance or require optimization of the ligation root for efficient addition of the first barcode digit. In order to fully realize BALI’s potential, further work will likely be required to optimize ligation rates at each serial step, and/or to introduce strategies to minimise the impact of failed ligation events, such as the adoption of encoding schemes with error correction features to ‘rescue’ partial spatial barcodes^50^.

A major limitation of state-of-the-art spatial multi-omics methods is their reliance on bespoke equipment that is inaccessible to most labs. By contrast, BALI has two main requirements: 1) prompt tissue segmentation and barcode assignment prior to the barcode writing; and 2) precise patterning of light from a single source to parallelise barcode assignment to all the areas with the same digit *i* in position *n.* While tissue segmentation and barcode assignment are computationally intensive processes, the wider access to high-performance graphic cards as part of the AI revolution has turned this seemingly limiting step into a routine process that can be completed within minutes. And while laser scanning optical setups – such as a confocal microscope – are not well suited to simultaneous illumination of multiple tissue segments, digital micromirror devices (DMDs) offer the opportunity to precisely pattern the light emitted from a light source, thus illuminating all the regions of interest at once. Therefore, we believe that BALI has the potential to provide a significant improvement in accessibility over other spatial omics technologies.

Nucleic acids are the natural targets of any method based on a sequencing readout. However, there is the potential to profile a broader portfolio of biomolecules by incorporating oligonucleotide-functionalized libraries of antibodies ^51, 26^. Antibodies can detect a broad spectrum of biomolecules (e.g. proteins, post-translational modifications, metabolites, lipids), yet they often suffer from poor specificity and limited resilience to oligonucleotide functionalisation. We envision that the expanding catalogue of single chain antibodies^52^ and the development of orthogonal chemistries to specifically decorate target molecules with oligonucleotides (e.g. aptamers^53^) will significantly extend the scope of spatial multi-omic profiling abilities in future work. Together with recent advances in obtaining high-quality transcriptomics profiles from FFPE tissue samples^54,55^, we believe that complementary innovations on multiple fronts are now coalescing to expand the applicability of spatial multi-omic technologies to new sample types and readouts – all of which should be compatible with BALI.

Considered together, BALI conceptually overcomes many limitations in the spatial multi-omics field, providing a high-resolution, adaptable, and accessible platform for optical, histology-informed barcoding that is poised to capitalize on emerging technical advances to further increase the scope, resolution, and applicability of spatial multi-omics profiling.

## Supporting information

Consortium Author List

## Acknowledgements

We would like to thank the members of the Hannon, Balasubramanian, and IMAXT consortium at the CRUK-Cambridge Institute and Department of Chemistry for their support and helpful discussions. We would also like to thank Lauren Deighton, Isabella Pearsall and Martin Tam with experimental support in the early stages of the project, and David Tannahill for experimental troubleshooting. A special acknowledgement to Clare Rebbeck for support with animal licencing and reporting, and the Hannon lab admin and lab management teams for their priceless help, in particular that of Heather Ashmore, Erica MacKenzie, and Michele Dunn. This work was enabled by the support received from the Lab Management, Biological Research Unit, Research Instrumentation, Microscopy and Genomics Cores at CRUK Cambridge Institute, with a special mention to Johanna Barbieri for her prompt assistance and useful advice on sequencing and Cathy Pauley for assisting with the logistics of light-protected experimental conditions. Additionally, we are indebted to Yuqi Chen, Angie Kirchner and Sean Flynn from the Balasubramanian group for giving us access and support for additional MiSeq and NextSeq runs. We would like to express our gratitude to April Pawluk from Life Science Editors for her invaluable assistance in editing the language and improving the narrative flow of this manuscript. This work was supported by the IMAXT Cancer Grand Challenge grant (A24042 to GJH and DB), CRUK Pioneer Award (G104344 to DB), Wellcome Trust Investigator Award (110161/Z/15/Z to GJH), a Cancer Research UK core award (A21143 to GJH) and Herchel Smith funds (to SB). GJH is a Royal Society Wolfson Research Professor (RSRP\R\200001) and Wellcome Trust Investigator. SB is a Herchel Smith Professor (University of Cambridge) and Wellcome Trust Senior Investigator (209441/Z/17.Z). ST-G was partially supported by The Branco Weiss Fellowship - Society in Science, administered by ETH Zürich. The authors’ work at the University of Cambridge is supported by the NIHR Cambridge Biomedical Research Centre (BRC-1215-20014). The views expressed are those of the authors and not necessarily those of the NIHR or the Department of Health and Social Care.

## Authors contributions

GB wrote this paper with contributions from DB, ST-G, BCN. and GJH. ST-G, GB and DB prepared the figures with input from BNC and GJH. GB, DB, CYS, EW, ST-G, SC, MA and NH performed the experiments relating to in situ ligation upon uncaging. CYS performed the overhang screening and validation assays. DB, GB and CB performed the experiments related to RNA-seq BALI. ST-G, GB, and SC performed the experiments related to ATAC-seq BALI. GB, ST-G, SC, and CB performed the experiments related to multi-omics BALI. DB performed the data analysis with input from GB and ST-G. DB, BCN, GJH, SB, GB, and ST-G contributed to discussions and study designs. DB and GH supervised the study.

## Data Availability

All the sequencing data and relative processed data pertinent to this work has been deposited in the Gene Expression Omnibus database with accession number GSE260699. Data will be released upon publication. Raw images and non-sequencing data are available from the authors upon request.

## Code Availability

No new algorithms or analysis tools were designed for this work. The scripts used for analysis and data processing are available from the authors upon request.

## Materials and Methods

### Overhang screen

Eight oligonucleotides (BL354 to BL357, and BL354B to BL357B) were synthesized with normal desalting purification. The oligos were annealed to produce two double-stranded DNA molecules (A and B) containing a constant region corresponding to the Truseq P5 and P7 sequences plus a multiplexing sequencing index, and a random 6-mer or 7-mer sequence protruding at the 5’ end (ligation overhang). Oligos were annealed at 30 μM concentration in 2X SSC by running the following program in a thermocycler: 1) 95°C for 5 minutes, 2) ramp down to 65 at 0.1°C /second, 3) hold at 65°C for 5 minutes, 4) ramp down to 12°C at 0.1°C/second, 5) hold at 12°C. The A and B dsDNA molecules for each overhang length (6-mer or 7-mer) were diluted to 3.334 μM in 2X SSC and used to assemble a ligation reaction as follows: 2 μL 10x T4 ligase buffer, 1 μL of each annealed dsDNA (final 166.7 nM), 1 μL T4 DNA ligase (NEB M0202M), 15 μL nuclease-free water. Ligation was performed for 3 minutes at 22°C, following by heat-inactivation at 65°C for 10 minutes. The ligated molecules were purified by running the whole reaction volume on a 2% TBE agarose gel. The gel was stained in 1:10000 SYBR-gold (Thermo Scientific) and the bands corresponding to the ligation products were purified using the Qiagen Qiaquick gel extraction kit using the supplier’s standard protocol. Library concentration was estimated by Tapestation (DNA5000 screentape) and qPCR (KAPA Illumina library quantification kit).

The fastq files containing raw sequencing reads were first filtered using the cutadapt package (v4.1) using the following options: -a AGATCGGAA -m [length of overhang] -M [length of overhang +1]. This extracted the sequence corresponding to each ligated 6-mer or 7-mer overhang in the library. The resulting files were processed through a custom python script counting the number of occurrences of each overhang, producing a final .csv file with overhang counts.

### Overhang screen validation

DNA indexes with different overhang sequences were validated by performing in-situ ligation assays on 4T1 mouse cells grown on 25mm round coverslips coated in poly-L-lysine overnight (Sigma P4707, 0.01% w/v). Cells were grown for 24-48hrs to ∼90% confluence.

Indices for each overhang to be tested, as well as the BL092_50nt root oligo, were ordered as oligonucleotides with a 5’ phosphate on the overhang subjected to ligation. For each overhang to be tested, two indices were annealed. BL407 to BL426 (6-mers) or BL452 to BL471 (7-mers) were annealed to BL427 to extend the root with the overhang ‘XX’ to be tested (‘Index_A_XX’), whereas BL428 to BL447 (6-mers) or BL472 to BL487 were annealed to BL448 to generate a reverse complement overhang to the one tested (‘Index_B_XX’). To increase phosphorylation yield, oligos with 5’ phosphate were further phosphorylated by incubating them (at a concentration of 50 μM for indices and 25 μM for BL092-50nt) with 10U T4 polynucleotide kinase (NEB) in 1X T4 DNA ligase buffer (NEB). The reaction was carried on for 30 minutes at 37°C and stopped by heating to 60°C for 20 minutes, followed by purification through Illustra Microspin G-25 columns (Cytiva) as per supplier’s protocol. DNA indices were annealed to a final concentration of 20 μM in 2X SSC using the following protocol in a thermal cycler: 1) 95°C for 5 minutes, 2) ramp down to 65 at 0.1°C /second, 3) hold at 65°C for 5 minutes, 4) ramp down to 12°C at 0.1°C /second, 5) hold at 12°C. Cells were fixed in 4% PFA/PBS for 15 minutes at RT, washed in PBS, and permeabilized in 1% Triton X-100 in PBS for 15 minutes at RT. After permeabilization, cells were further cleared and permeabilized by incubating them in 0.3% SDS in PBS for 30 minutes at RT, followed by three washes in PBS for 10 minutes each. Hybridization of the root was performed by inverting the coverslips on a drop (25 μl) of 0.1 μM polyT-rootV2 oligo diluted in hybridization buffer (2x SSC, 1 mg/mL yeast RNA, 10% dextran sulfate, 1:1000 Murine RNase inhibitor NEB M0314) on a parafilm sheet placed on a glass plate. Hybridization was performed for 24hrs at 37°C in a humidified chamber. The following day, coverslips were transferred to a 6-well plate and washed three times with 1 mL of 2X SSC for 5 minutes each and once in secondary hybridization buffer (2X SSC, 10% Ethylene Carbonate) for 5 minutes, all at RT. Samples were then hybridized with 2 ml of 1.25 nM BL092-50nt oligo in secondary hybridization buffer for 20 minutes at RT, followed by three more washes in 2X SSC for 5 minutes each.

A ligation reaction was then assembled by combining 2 μl 10x T4 DNA ligase buffer (NEB), 1 μl annealed 20 μM Index_A_XX stock, 16ul nuclease-free water and 1 μl T4 DNA ligase (NEB). The entire volume of the reaction was pipetted on a parafilm sheet placed on a glass plate and the coverslips were inverted over it. Ligation was carried out at 25°C for 45 minutes in a humid chamber. After the ligation, coverslips were washed 3 times in 2X SSC for 5 minutes each to remove any unligated indices. A second ligation was performed as above using the cognate Index_B_XX.

Ligated root molecules were eluted by pre-heating plastic surface (the lid of a 6-well multiwell plate) at 95°C on a thermoblock. 50 μl of pre-heated 2X RNA loading buffer (95% deionized formamide, 5% 10X TBE, 5mg bromophenol blue) were pipetted on the surface and the coverslips were inverted on them. Samples were incubated for 10 minutes at 95°C, after which the coverslips were carefully lifted, and the liquid collected under them collected with a 200um pipette (approx. 30 μl were recovered per sample) and transferred to a 1.5ml Eppendorf tube. Samples were further incubated at 95°C for 10 minutes in a thermoblock and loaded on a 6% TBE-Urea acrylamide denaturing gel (Thermo Scientific). Gels were stained with SYBR-gold (1:10000 in 1x TBE, Thermo Scientific) and imaged using an Amersham Typhoon instrument in the cy2 and cy5 channels. Ligation efficiency was evaluated by densitometry on the cy5 gel images.

### Barcode writing on slide

Amine coated glass slides (Silane-Prep, Sigma, S4651) were washed in 70% Ethanol and air dried. All furthers steps were performed in light protected conditions. A functionalisation mix (50 μL) was prepared fresh with the 3’aminated 5’ caged oligo root carrying a terminal Cy3 dye (2 μM BL003 for the single-cycle experiment and 2 μM BL728 for the 4-cycle experiment) and a bifunctional crosslinker BS(5)PEG (Thermo Scientific) (5 μM in dimethylformamide), and placed at the centre of the slide as a drop. The functionalization was run 16 hrs at 37C in a humidified chamber. The functionalised slides were washed twice in 2X SSC (Thermo Scientific), and then mounted with 200 μL of 2x SCC and a thin glass coverslip (22×22mm).

For the single-cycle experiment, uncaging was performed on a SP5 Leica confocal system fitted with a 30mW 405nm solid state laser, illuminating for 2 minutes (“1” shape) and 5 minutes (“2” shape) with the 405nm laser at maximum power through a 10X objective. After uncaging the slide was imaged in the cy3 channel using the 514nm laser line to detect the removal of the fluorophore in the uncaged areas. The slide was then washed 3 times for 5 minutes each in 2X SSC. Oligos *lig_test_onbridge1_fw\** and *lig_test_onbridge1_rev* were annealed by mixing them at 45 μM final concentration in 2X SSC and running the following program in a thermocycler: 1) 95°C for 5 minutes, 2) ramp down to 65 at 0.1C/second, 3) hold at 65C for 5 minutes, 4) ramp down to 12C at 0.1C/second, 5) hold at 12°C. A ligation mix was then prepared with the annealed index oligos (2.7 μM), T4 ligase (100 U/μL, NEB) in 1x DTT free ligation buffer in a final volume of 50 μL. Any excess liquid from previous washes was removed from the slides, and 50uL of ligation mix were added to the marked uncaged area, covered with a coverslip and incubated for 60 minutes at 25°C in a humid chamber. After ligation, the slide was washed in 2X SSC for five minutes three times, in 0.2X SSC for five minutes once, and then in 0.2X SSC overnight. The following day the slide was imaged in the cy3 and cy5 channels using the 514nm and 633nm laser lines to detect the cy5-positive ligation products.

For the 4-cycle experiment, uncaging was performed as above illuminating for 15 minutes with the 405nm laser at maximum power. The sixteen barcode areas were first defined as separate square ROIs, then selected appropriately for each uncaging cycle. After uncaging, the slides were imaged as above (Cy3 and Cy5 channels) and then washed for 5 minutes in 2x SSC, 5 minutes in 0.2x SSC, and 5 minutes in 1x DTT free ligation buffer (50 mM TrisHCl pH7.4, 10 mM MgCl2, 1 mM ATP). The index oligos for each ligation were annealed as above. The ligation mix was prepared with the annealed index oligos (2.7 μM), T4 ligase (100 U/μL, NEB) in 1x DTT free ligation buffer in a final volume of 50 μL. Any excess liquid from previous washes was removed from the slides, and 50 μL of ligation mix were added to the marked uncaged area, covered with a coverslip, and incubated for 60 minutes at 25°C in a humid chamber. After ligation, the slides were washed 10 minutes in 2x SSC, 5 minutes in 2x SSC + 10% Ethylene Carbonate, 5 minutes in 2x SSC, then mounted with 200 μL of 2x SSC for the additional round of uncaging.

### Multiple ligations on beads

Sepharose NHS beads (50 μL of resin bed) were washed twice in 20 volumes of 1 mM HCl, and twice in 20 volumes of 2x Coupling Buffer (100 mM Sodium Borate pH 8.5). After the last wash, the resin bed was resuspended in 2 volumes of 50 μM of 5’ amino modified oligo (PolyT_root_probe_v2). The beads were functionalised for 4 hrs at room temperature with constant rotation. The reaction was blocked by adding 20 μL of 1M Tris-HCl pH 8 and rotating the beads for additional 30 minutes at room temperature. The beads were washed four times for 5 minutes in 100 mM Tris-HCl pH 8 and either used immediately or stored in 1x Storage buffer (100 mM Tris-HCl pH 8, 2.5 mM EDTA) for up to 1 month at 4°C. Prior to the ligations, all oligos purchased with a 5’ phosphate were additionally enzymatically phosphorylated with 0.2 U/μL T4 PNK, 1 mM ATP, 0.5 μM 5’P oligo in 1x PNK reaction buffer for 30 minutes at 37°C, and then heat-inactivated for 20 minutes for 60°C. Samples corresponding to a different number of serial ligation cycles were processed in parallel, removing one sample at each cycle for analysis. For each sample, 12.5 μL of functionalised bead resin were washed twice in 1x Hybridization buffer (2x SSC, 10% Ethylene Carbonate in H20) and hybridised with 100 μL of a 3’ Cy5 labelled 5’P oligo root (BL92) (1 μM in 1x Hybridization buffer) for 30 minutes at room temperature with constant rotation. To remove any free fluorescent oligo, the beads were washed 3 times for 10 minutes in 1x Hybridization buffer, and 2 times for 5 minutes in 2x SSC with constant rotation. The index oligos for the ligations were annealed at 45 μM final in 2x SCC in the following pairs: BL46-PNK/BL45 (cycles 1, 3, 5, 7) and BL47-PNK/BL48 (cycles 2, 4, 6). The annealing thermocycler conditions were as follows: 95°C for 5 minutes, ramp down to 65°C at 0.1°C /sec, 65°C for 5 minutes, ramp down to 4°C at 0.1°C /sec, hold at 4°C. For each cycle of ligation, a ligation mix (20 μL) was prepared with annealed index oligos (2.7 μM) alternating BL46/45 and BL48/47, T4 ligase (100U/μL, NEB) in 1x T4 ligation buffer in a final volume of 20 μL and added directly to the beads. The ligation reactions were incubated for 30 minutes at room temperature with constant rotation, and then washed 3 times for 5 minutes in 2x SSC. After the serial ligations, the ligated products were denatured from the beads by boiling the beads at 95°C for 5 minutes in 1x RNA loading dye (47.5% formamide, 0.01% SDS, 0.01% bromophenol blue, 0.5 mM EDTA). The beads were centrifuged and the supernatant run on a 15% TBE-Urea gel. Images of the gels were acquired with an Amersham Typhoon.

### Animal housing

All animal procedures were performed in accordance with the Animal (Scientific Procedures) Act 1986 under UK Home Office project license PAD85403A. Animals were housed under standard husbandry conditions per UK Home Office and Institute regulations. Individually ventilated cages housed four to five animals under a 12-h light, 12-h dark cycle at the CRUK Cambridge Institute animal facility. Relative humidity was kept between 45 and 65% with a temperature range of 20-24°C. Animals had free access to water and pelleted food. Embryos samples were taken at embryonic day 16.5 from ethically euthanised pregnant dams following timed mating. Adult samples were collected from ethically euthanised 8 −12 month-old animals.

### Tissue collection and Histology

For embryo collection, pregnant dams were euthanised by CO2 overdose with exposure to gas in rising concentration followed by cervical dislocation. Embryos were euthanised by cooling on ice followed by decapitation. Heads were frozen in OCT resin by immersion in a dry ice/isopentane bath followed by storage at −80°C. For adult brain collection, mice were euthanised by cervical dislocation followed by confirmation of death by permanent cessation of circulation. Animals were then decapitated, and the brain dissected and frozen in OCT resin and stored as above. Cryosections were cut using a Leica CM3050S cryostat under RNAse free conditions. The instrument was wiped with RNAse-ZAP prior to use and between different blocks. A different blade was used for each block. Sample blocks were equilibrated to the chamber temperature for 15 minutes prior to sectioning, and then mounted on the object holder. Sections were cut at a thickness of 10 μM and collected on RNAse-free SuperFrost Plus microscope slides (one section per slide). Slides were left at room temperature for 20-30 seconds to help tissue adhesion and then stored on dry ice (short term) or at −80°C (long term). Sections to be processed in the Laser-Capture microdissection workflow were collected on PEN membrane glass slides (Leica) and processed/stored similarly.

### Multiple ligations on tissue

All steps were performed under RNAse-free conditions. Cryosections of adult liver were retrieved from cold storage and equilibrated for 5 minutes at room temperature and then heated for 5 minutes at 37C on a metal heat block. Pap-pen was used to draw an hydrophobic boundary around each section, and let dry for 5 minutes at room temperature. Sections were incubated with 500 μL PBS to dissolve and remove the OCT embedding matrix and fixed with 500 μL of 0.2% PFA in PBS for exactly 5 minutes at room temperature. Fixation was quenched by removing the PFA and promptly adding 1.25 M Glycine in PBS, one quick wash and one wash for 5 minutes at room temperature. After washing twice in 1x wash buffer (20 mM HEPES pH 7.5, 150 mM NaCl, 0.5 mM spermidine, 1x proteinase inhibitor cocktail) for 5 minutes each, the sections were permeabilised in Permeabilization buffer (0.01% IGEPAL, 0.01% Digitonin in 1x Wash buffer) for 5 minutes at room temp, and then washed in 1x Wash buffer twice for 5 minutes. The permeabilised section were heated for 5 minutes in pre-warmed 1x PBS at 65°C in the hybridization oven, and then snap-cooled in ice-cold PBS on ice for 10 minutes. For each slide, a mock RT mix was prepared as follows to hybridise a root for sequential ligations. A first mix (14 μL) containing 2.5 μL of dNTPs (stock 10 mM each), 1.5uL of poly-T root (100 μM stock) in water was denatured for 5 minutes in 65°C and the snap cooled on ice for >2 minutes. A second mix (36 μL) was prepared with 10 μL of 5x SuperScript-IV buffer, 1 μL of RNAsin, 2.5 μL of 100 mM DTT in nuclease free water and added to the first mix. Any excess PBS was removed from the slides, and 50 μL of mock RT mix was added directly on top of the sections, which were then covered with a 22×22 mm coverslip and incubated overnight (∼16hrs) at 42°C in a sealed humid chamber. Prior to the ligations, all oligos purchased with a 5’ phosphate (BL611, BL604 and BL603) were additionally enzymatically phosphorylated with T4 PNK (1.6 U/μL), ATP (1 mM), 5’P oligo (50 μM) in 1x T4 reaction buffer for 60 minutes at 37°C and then heat-inactivated for 20 minutes for 60°C. The oligos were purified through a G25 resin column following the manufacturer standard protocol. The index oligos for the ligations were annealed at 30 μM final in 2x SCC in the following pairs: BL604-PNK/BL585 (cycles 1 and 3) and BL603-PNK/BL583 (cycle 2). The annealing thermocycler conditions were as follows: 95°C for 5 minutes, ramp down to 65°C at 0.1°C /sec, 65°C for 5 minutes, ramp down to 4°C at 0.1°C /sec, hold at 4°C. The slides were retrieved from the mock RT reaction, and washed twice in PBS, once in 2x SSC and once in 2x SSC + 10% EC for 5 minutes each at room temperature. The fluorescent ligation root was hybridised to the poly-T root by incubating the sections with 500 μL of 12.5nM BL611-PNK in 2x SSC + 10%EC for 30 minutes at room temperature. The sections were washed 3 times in 2x SSC and used for serial ligations. For each number of cycles, we processed two slides in parallel as replicates. For each cycle of ligation, a ligation mix (25 μL) was prepared with annealed index oligos (3 μM, alternating BL604/585 and BL603/583 as appropriate), T4 ligase (100 U/μL, NEB) in 1x T4 ligation buffer, added directly to the sections and covered with a 22×22mm glass coverslip. The ligation reactions were incubated for 30 minutes at 25°C in a humid chamber, and then washed 3 times for 5 minutes in 2x SSC. After the serial ligations, the ligated products were denatured from the sections by boiling the slides at 95°C for 15 minutes in 100uL of 1x RNA loading dye (47.5% formamide, 0.01% SDS, 0.01% bromophenol blue, 0.5 mM EDTA) on a flat heat block under humid conditions. The RNA loading dye was retrieved (∼80uL) in a PCR tube, additionally denatured for 10 minutes at 95°C and run on a 8% TBE-Urea gel. Images of the gels were acquired with an Amersham Typhoon.

### Laser Capture microdissection on E16.5 mouse brain

Tissue sections from E16.5 mouse embryos (as described in the “Animal housing” section) were cut as described in the “Tissue collection” section. Sections were stained with Haematoxylin/Eosin using the following fast procedure: 1) 40 seconds in 75% ethanol, 2) 30 seconds in mQ water, 3) 30 seconds in Harris’ modified Haematoxylin solution (Sigma HHS16), 4) 3x 30 seconds in mQ water, 5) 30 seconds in a 1:1000 dilution of 28% Ammonium Hydroxide (Sigma 338818) in mQ water, 6) 10 seconds in 5% Eosin Y solution (Sigma 318906), 7) 30 seconds each in 70%, 95% and 100% ethanol, 7) 2x 20 seconds in Xylene. The stained sections were imaged and cut on a Leica LMD6000 LCM microscope. Areas corresponding to the sub-ventricular zone and to the cortex (matching the illumination areas for the BALI experiment) were used for cutting. Tissue fragments were collected in the caps of 0.5ml RNAse-free tubes in 50 μl of lysis solution from the RNAqueous micro RNA extraction kit (Thermo Scientific). RNA extraction was performed according to the RNAqeous micro kit protocol for LCM extraction. In short: 1) 50 μl of lysis buffer were added to the 50 μl including the sample, and the tube was incubated at 42°C for 30 minutes, 2) the collection column was pre-wetted with 30 μl lysis solution, 3) 3 μl LCM additive were added to each sample, 4) 129 μl of 100 μl Ethanol were added to each sample, 5) the samples were bound to the collection column, and the column was washed with 180 μl wash solution 1 and twice with 180 μl of wash solution 2/3, 6) RNA was eluted with a 5 minute incubation step with 10 μl of pre-heated (95°C) elution solution. Purified RNA was treated with DNAse using the optional DNAse treatment step in the RNAqueous kit: 1) 1 μL of DNAseI buffer and 1 μL of DNAse enzyme were added to each sample, 2) samples were incubated for 20 minutes at 37°C, 3) 2 μL of DNAse inactivation reagent were added to each sample, and the inactivated DNAseI was removed by centrifugation. RNA abundance and quality were evaluated by Agilent Tapestation (DNA5000 High-Sensitivity screentape). RNA sequencing libraries were prepared using the NEBNext Single Cell/Low Input RNA Library Prep Kit for Illumina (NEB) according to the recommended protocol using 1 ng RNA as input material. Library quality and abundance was measured by Tapestation (DNA5000 screentape) and qPCR (KAPA Illumina library quantification kit). Libraries were sequenced on a NextSeq500 instrument using paired end 75nt reads.

### In situ RNA-seq

All steps were performed under RNAse-free conditions. Cryosections of embryonic E16.5 brain (coronal plane) were retrieved from cold storage and equilibrated for 5 minutes at room temperature and then heated for 5 minutes at 37°C on a metal heat block. Pap-pen was used to draw a hydrophobic boundary around each section and let dry for 5 minutes at room temperature. Sections were incubated with 500 μL PBS to dissolve and remove the OCT embedding matrix and fixed with 500 μL of 0.5% PFA in PBS for exactly 5 minutes at room temperature. Fixation was quenched by removing the PFA and promptly adding 1.25M Glycine in PBS, one quick wash and one wash for 5 minutes at room temperature. After washing twice in PBS for 5 minutes each, the sections were permeabilised in Permeabilization buffer (0.5% TritonX100 in PBS) for 15 minutes at room temp, and then washed in PBS 3 times for 5 minutes. The sections were incubated with 0.1N HCl for 5 minutes and washed 3 times in PBS (two quick washes, one 5 minutes wash). The section were heated for 5 minutes in pre-warmed PBS at 65°C in an hybridization oven, and then snap-cooled in ice-cold PBS on ice for 10 minutes. From this point onwards all steps performed in light protected conditions. For each slide, an first RT mix (14uL) was prepared containing 2.5 μL of dNTPs (stock 10 mM each), 1.5μL of caged RT primer (BL621, 100 μM stock) in water, denatured for 5 minutes in 65C and the snap cooled on ice for >2 minutes. A second RT mix (36 μL) was prepared with 10 μL of 5x SuperScript-IV buffer, 2.5 μL SuperScript-IV enzyme, 1 μL of RNAsin, 3 μL of 100 μM TSO oligo (BL617), 5 μL of 5M Betaine, 0.5 μL ET-SSB, 2.5 μL of 100 mM DTT in nuclease free water, and added to the first mix, up to a total RT reaction volume of 50uL. Any excess PBS was removed from the slides, and 50 μL of RT mix was added directly on top of the sections, which were then covered with a 22×22 mm coverslip and incubated overnight (∼16hrs) at 42°C in a sealed humid chamber. The index oligos for the ligations were annealed at 45uM final in 2x SCC in the following pairs: BL599/BL623 (cycle 1, SZV) and BL601/BL624 (cycle 1, cortex). The annealing thermocycler conditions were as follows: 95°C for 5 minutes, ramp down to 65°C at 0.1°C /sec, 65°C for 5 minutes, ramp down to 4°C at 0.1°C /sec, hold at 4°C. The slides were retrieved from the RT reaction, washed one in PBS then twice in 2x SSC for 5 minutes each. Nuclei were stained by incubating the section with Draq5 (1:1000 dilution in 2x SSC) for 10 minutes, and then washed 2 times for 5 minutes in 2x SSC. Since the RT primer was directly caged, we proceed with two consecutive rounds of uncaging/ligation to mark the SVZ and cortex areas (bilaterally). For the uncaging, the slides were mounted with 200 μL of 2x SSC and a thin glass coverslip (22×22mm). The uncaging was performed on a SP5 Leica confocal system fitted with a 30mW 405nm solid state laser, illuminating each ROI for 5 minutes with the 405nm laser at maximum power. The slides were washed in 2x SSC, once quickly and once for 5 minutes, and then incubated for 5 minutes in 1x Ligation buffer (50 mM TrisHCl pH7.4, 10 mM MgCl2, 10 mM DTT, 1mM ATP). For each sample, a ligation mix (50 μL) was prepared as follows: index oligos (3 μM, alternating BL599/BL623and BL624/BL601 as appropriate), T4 ligase (100U/ μL, NEB) in 1x T4 ligation buffer. The ligation mix was added directly to the sections, covered with a 22×22mm glass coverslip, and the reactions were incubated for 30 minutes at 25°C in a humid chamber. The slides were then washed in 2x SSC + Draq5 (1:1000) once for 10 minutes, 2x SSC + 10%EC once for 5 minutes and 2x SSC once for 5 minutes. After the serial ligations, a hybridization chamber was mounted on the slides, and the sections were lysed in 200 μL of Lysis Buffer (0.8 U/mL in RIPA buffer) at 60°C overnight (∼16 hrs) in a sealed humid chamber. All the following steps were not light protected anymore. The lysate was collected and purified with MinElute Reaction Clean-up column (Qiagen), and eluted in 25 μL of EB buffer. A small aliquot of the lysate (2 μL) was used as input for a qPCR to measure the optimal cycling conditions for the amplification of the libraries. The qPCR reaction was set up as follows: 2 μL of purified lysate, 25 μL of 2x Q5 master mix, 0.25 μL of 50 μM universal forward Truseq primer, 0.5 μL of 50 μM index A012 Truseq primer 2.5 μL of 20x EVA-Green. The cycling protocol was 98°C for 3 minutes, 40 cycles of 98°C for 15 seconds, 68°C for 20 seconds, 72°C for 20 seconds, plate reading step. The number of cycles required to reach 1/3 of the maximum signal was selected for the final amplification of the libraries. The final amplification reaction was set as follows: 10 μL of purified lysate, 25 μL of 2x Q5 master mix, 0.25 μL of 50 μM universal forward Truseq primer, 0.5 μL of 50 μM index A0XX Truseq primer, 2.5 μL of 20x EVA-Green. The cycling conditions were the same as the qPCR. The PCR reactions were purified with AmpureXP beads (0.8x) and eluted in 20 μL of EB buffer. The libraries were analysed on a Tapestation (D5000 cartridge) and quantified with KAPA library quantification kit prior to pooling. Sequencing was performed on a Novaseq (Illumina) instrument.

### Transposome assembling

The adapters for transposome assembling were annealed at 50 μM each in 1x Annealing buffer (40 mM TrisHCl pH 8.0, 50 mM NaCl) with the following thermocycling protocol: 95°C for 5 minutes, ramp down to 65°C at 0.1°C /sec, 65°C for 5 minutes, ramp down to 4°C at 0.1°C /sec, hold at 4°C. The oligos pairs used were BL699/BL538 (BALI-adapter, universal ligation root) and BL515/BL538 (A-adapter, single modality ATAC) or BL749/BL538 (A-adapter, multi-omics). The transposomes were assembled by mixing 3 μL of annealed BALI-adapter, 3 μL of annealed A-adapter, 6 μL of unloaded Tn5 and incubating the mix at 23°C for 30 minutes and then used immediately afterwards.

### In situ ATAC

Cryosections of adult brain were retrieved from cold storage and equilibrated for 5 minutes at room temperature and then heated for 5 minutes at 37°C on a metal heat block. Pap-pen was used to draw a hydrophobic boundary around each section, and let dry for 5 minutes at room temperature. Sections were incubated with 500 μL PBS to dissolve and remove the OCT embedding matrix, and fixed with 500 μL of 0.5% PFA in PBS for exactly 5 minutes at room temperature. Fixation was quenched by removing the PFA and promptly adding 1.25M Glycine in PBS, one quick wash and one wash for 5 minutes at room temperature. After washing twice in PBS for 5 minutes each, the sections were permeabilised in Permeabilization buffer (10 mM TrisHCl pH 7.4, 10 mM NaCl, 3 mM MgCl_2_, 0.01% Tween-20, 0.01% IGEPAL, 0.001% Digitonin, 1% BSA, 10 μL murine RNAse Inhibitor) for 15 minutes at room temperature. The sections were washed in 1x Wash Buffer (10 mM TrisHCl pH 7.4, 10 mM NaCl, 3 mM MgCl_2_, 0.1% Tween-20, 1% BSA, 10 μL murine RNAse Inhibitor) once quickly and once for 5 minutes, then in PBS twice for 5 minutes. For each section, 200 μL of transposition mix were prepared as follows: 2 μL of loaded Tn5 in 10 mM Tris-HCl pH 7.6, 5 mM MgCl2, 10% Dimethyl Formamide, 0.33x PBS, 0.1% Tween-20, 0.01% Digitonin. Each section was sealed with an adhesive hybridization chamber and incubated with 200 μL of tagmentation mix for 35 minutes at 37°C in a humid chamber. The slides were retrieved and washed with 40 mM EDTA once quickly, and once for 5 minutes, then 3 times in 1x PBS for 5 minutes each and 2 times in 2x SSC for 5 minutes each. The index oligos for the ligations were annealed at 45 μM final in 1xSSC in the following pairs: BL680/BL676 (cycle 1, bulk), BL681/BL677 (cycle 2, bulk), BL683/BL642 (cycle 3, bulk), BL732/BL601 (cycle 4A, Dentate Gyrus), BL733/BL677 (cycle 4B, cortex), BL732/BL641 (cycle 1, single ligation bulk). The annealing thermocycler conditions were as follows: 95°C for 5 minutes, ramp down to 65°C at 0.1C/sec, 65°C for 5 minutes, ramp down to 4°C at 0.1°C /sec, hold at 4°C. The spatial barcodes were encoded in the tissue with the following protocol, which was conducted under light protected condition for its entirety: cycle 1 ligation, cycle 2 uncaging (bulk), cycle 2 ligation, cycle 3 uncaging (bulk), cycle 3 ligation, cycle 4A uncaging (Dentate Gyrus), cycle 4A ligation, cycle 4B uncaging (cortex), cycle 4B ligation. Bulk uncaging was performed with a UV transilluminator fitted with 365 nm bulbs for 15 minutes at room temperature, with the section covered with 500 μL of 2x SSC. Spatial uncaging was performed on a SP5 Leica confocal system fitted with a 30mW 405nm solid state laser, illuminating each ROI for 10 minutes with the 405nm laser at maximum power. Following uncaging, the slides were washed in 2x SSC, once quickly and once for 5 minutes. Ligations were performed as follows. Sections were incubated for 5 minutes in 1x Ligation buffer (50 mM TrisHCl pH7.4, 10 mM MgCl2, 10 mM DTT, 1mM ATP). For each sample, a ligation mix (50 μL) was prepared as follows: index oligos (3 μM), T4 ligase (100 U/μL, NEB) in 1x T4 ligation buffer. The ligation mix was added directly to the sections, covered with a 22×22mm glass coverslip, and the reactions were incubated for 30 minutes (45 minutes for cycle 1) at 25°C in a humid chamber. The slides were then washed in 2x SSC + Draq5 (1:1000) once for 10 minutes, 2x SSC + 10%EC once for 5 minutes and 2x SSC once for 5 minutes. After the serial ligations, an adhesive hybridization chamber was mounted on the slides, and the sections were lysed in 200 μL of Lysis Buffer (0.8 U/mL in RIPA buffer) at 60°C overnight (∼16 hrs) in a sealed humid chamber. All the following steps were not carried out under light protected conditions. The lysate was collected and purified with MinElute Reaction Clean-up column, and eluted in 25 μL of EB buffer. Libraries were amplified via quantitative PCR as follows. For each sample, a 50 μL PCR mix was prepared mixing 5 μL of purified sample, 25 μL of 2x Q5 Mastermix, 1.25 μL of 50 μM P5 primer (BL727), 1.25 μL of 50 μM indexed P7 primer (TruSeq A0XX) in nuclease free water. The reaction was incubated in a thermocycler at 72°C for 5 minutes (gap-fill), 98°C for 3 minutes, then 5 cycles at 98°C for 20 seconds, 68°C for 20 seconds, 72°C for 20 seconds. The pre-amplified products was kept on ice, and 5 μL were used as input for qPCR to determine the optimal number of additional cycles, The qPCR reactions were set up as follows: 5 μL of the pre-amplified material, 5 μL of 2x Q5 master mix, 0.25 μL of 50 μM P5 primer (BL727), 1.25 μL of 50 μM indexed P7 primer (TruSeq A0XX), 2.5 μL of 20x EVA-Green in nuclease free water. The cycling protocol was 98°C for 3 minutes, 40 cycles of 98°C for 20 seconds, 68°C for 20 seconds, 72°C for 20 seconds with a plate reading step. The number of cycles required to reach 1/3 of the amplification curve plateau was selected for the final amplification of the libraries. The PCR reactions were purified with AmpureXP beads (0.8x) and eluted in 20 μL of EB buffer. The libraries were analysed on a Tapestation (D5000 cartridge) and quantified with KAPA library quantification kit prior to pooling. Sequencing was performed on a Novaseq (Illumina) instrument.

### In situ multi-omics

All steps were performed under RNAse-free conditions. Cryosections of adult brain were retrieved from cold storage and equilibrated for 5 minutes at room temperature and then heated for 5 minutes at 37°C on a metal heat block. Pap-pen was used to draw a hydrophobic boundary around each section, and let dry for 5 minutes at room temperature. Sections were incubated with 500 μL PBS to dissolve and remove the OCT embedding matrix, and fixed with 500 μL of 0.5% PFA in PBS for exactly 5 minutes at room temperature. Fixation was quenched by removing the PFA and promptly adding 1.25 M Glycine in PBS, one quick wash and one wash for 5 minutes at room temperature. After washing twice in PBS for 5 minutes each, the sections were permeabilised in Permeabilization buffer (10 mM TrisHCl pH 7.4, 10 mM NaCl, 3 mM MgCl_2_, 0.01% Tween-20, 0.01% IGEPAL, 0.001% Digitonin, 1% BSA, 10 μL murine RNAse Inhibitor) for 15 minutes at room temp. The sections were washed in 1x Wash Buffer (10 mM TrisHCl pH 7.4, 10 mM NaCl, 3 mM MgCl_2_, 0.1% Tween-20, 1% BSA, 10 μL murine RNAse Inhibitor) once quickly and once for 5 minutes, then in PBS twice for 5 minutes. For each section, 200 μL of transposition mix were prepared as follows: 2uL of loaded Tn5 in 10 mM Tris-HCl pH 7.6, 5 mM MgCl2, 10% Dimethyl Formamide, 0.33x PBS, 0.1% Tween-20, 0.01% Digitonin. Each section was sealed with an adhesive hybridization chamber and incubated with 200 μL of tagmentation mix for 35 minutes at 37°C in a humid chamber. The slides were retrieved and washed with 40 mM EDTA once quickly, and once for 5 minutes, then with PBS for 3 times for 5 minutes. The sections were heated for 5 minutes in pre-warmed PBS at 65°C in a hybridization oven, and then snap-cooled in ice-cold PBS on ice for 10 minutes. For each slide, a first RT mix (14uL) was prepared containing 2.5 μL of dNTPs (stock 10 mM each), 1.5μL of phosphorylated RT primer (BL730, 100 μM stock) in water, denatured for 5 minutes in 65°C and the snap cooled on ice for >2 minutes. A second RT mix (36 μL) was prepared with 10 μL of 5x SuperScript-IV buffer, 2.5μL SuperScript-IV enzyme, 1 μL of RNAsin, 3 μL of 100 μM TSO oligo (BL617), 5 μL of 5M Betaine, 2.5 μL of 100 mM DTT in nuclease free water, and added to the first mix, up to a total RT reaction volume of 50 μL. Any excess PBS was removed from the slides, and 50 μL of RT mix was added directly on top of the sections, which were then covered with a 22×22 mm coverslip and incubated overnight (∼16hrs) at 42°C in a sealed humid chamber. The slides were retrieved from the RT reaction, washed one in 1x PBS then twice in 2x SSC for 5 minutes each. The index oligos for the ligations were annealed at 45 μM final in 1xSSC in the following pairs: BL680/BL676(cycle 1, bulk), BL681/BL677 (cycle 2, bulk), BL683/BL642 (cycle 3, bulk), BL732/BL601 (cycle 4A, Dentate Gyrus), BL733/BL677 (cycle 4B, CA1), BL732/BL641 (cycle 1, single ligation bulk). The annealing thermocycler conditions were as follows: 95°C for 5 minutes, ramp down to 65°C at 0.1°C /sec, 65°C for 5 minutes, ramp down to 4°C at 0.1°C /sec, hold at 4°C. The spatial barcodes were encoded in the tissue with the following protocol, which was conducted under light protected condition for its entirety: cycle 1 ligation, cycle 2 uncaging (bulk), cycle 2 ligation, cycle 3 uncaging (bulk), cycle 3 ligation, cycle 4A uncaging (Dentate Gyrus), cycle 4A ligation, cycle 4B uncaging (CA1), cycle 4B ligation. Bulk uncaging was performed with a UV transilluminator fitted with 365 nm bulbs for 15 minutes at room temperature, with the section covered with 500 μL of 2x SSC. Spatial uncaging was performed on a SP5 Leica confocal system fitted with a 30mW 405nm solid state laser, illuminating each ROI for 10 minutes with the 405nm laser at maximum power. Following uncaging, the slides were washed in 2x SSC, once quickly and once for 5 minutes. Ligations were performed as follows. Sections were incubated for 5 minutes in 1x Ligation buffer (50 mM TrisHCl pH7.4, 10 mM MgCl2, 10 mM DTT, 1mM ATP). For each sample, a ligation mix (50 μL) was prepared as follows: index oligos (3 μM, as appropriate), T4 ligase (100 U/μL, NEB) in 1x T4 ligation buffer. The ligation mix was added directly to the sections, covered with a 22×22mm glass coverslip, and the reactions were incubated for 30 minutes (45 minutes for cycle 1) at 25°C in a humid chamber. The slides were then washed in 2x SSC + Draq5 (1:1000) once for 10 minutes, 2x SSC + 10%EC once for 5 minutes and 2x SSC once for 5 minutes. After the serial ligations, an adhesive hybridization chamber was mounted on the slides, and the sections were lysed in 200 μL of Lysis Buffer (0.8 U/mL in RIPA buffer) at 60°C overnight (∼16 hrs) in a sealed humid chamber. All the following steps were not carried out under light protected conditions. The lysate (200 μL) was collected and an aliquot (150 μL) was treated with 10 μL of 100 mM PMSF for 10 minutes at room temperature to inactivate the proteinase K prior to biotin pulldown to separate the Tn5 fragments (biotinylated) from cDNA. For each sample, 10uL of MyOne C1 beads were washes 3 times in 1xB&W-T buffer, and then resuspended in 150 μL of 2xB&W-T buffer. The lysate was then added and incubated with the beads for 60 minutes at room temperature. After this Incubation we collected the supernatant as our cDNA enriched fraction for RNA libraries, while the beads represented our Tn5 fragments enriched fraction for ATAC libraries. The beads were washed 3 times in 1xB&W-T buffer for 5 minutes, once in 1xST buffer for 5 minutes and then directly used as input for library amplification via precise PCR as follows. For each sample, a 50 μL PCR mix was prepared mixing 25 μL of 2x Q5 Mastermix, 2.5 μL of 10 μM P5 primer (BL727), 2.5 μL of 10 μM indexed P7 primer (TruSeq A0XX) in nuclease free water, and adding it directly to the washed bead pellet. The reaction was incubated in a thermocycler at 72°C for 5 minutes (gap-fill), 98°C for 3 minutes, then 5 cycles at 98°C for 20 seconds, 68°C for 20 seconds, 72°C for 20 seconds. The pre-amplified products was kept on ice, and 5 μL were used as input for qPCR to determine the optimal number of additional cycles, The qPCR reactions were set up as follows: 5 μL of the pre-amplified material (after removal of the beads), 5 μL of 2x Q5 master mix, 0.5 μL of 10 μM P5 primer (BL727), 0.5 μL of 10 μM indexed P7 primer (TruSeq A0XX), 2.5 μL of 20x EVA-Green in nuclease free water. The cycling protocol was 98°C for 3 minutes, 40 cycles of 98°C for 20 seconds, 68°C for 20 seconds, 72°C for 20 seconds with a plate reading step. The number of cycles required to reach 1/3 of the amplification curve plateau was selected for the final amplification of the libraries. The PCR reactions were purified with AmpureXP beads (1.2x) and eluted in 20 μL of EB buffer. The libraries were analysed on a Tapestation (D5000 cartridge) and quantified with KAPA library quantification kit prior to pooling. Sequencing was performed on a Novaseq (Illumina) instrument. Separately, the cDNA fraction was purified using Monarch Reaction clean-up columns and eluted in 12.5uL. The cDNA was treated with RNAseH by adding 1.5 μL of 10x RNAseH buffer and 1 μL RNAseH enzyme to each sample, and then incubating the mix at 37°C for 30 minutes, 65°C for 20 minutes. The cDNA was amplified to via precise PCR as follows. For each sample, a 50 μL PCR mix was prepared mixing 15 μL of RNAseH treated sample, 25 μL of 2x Q5 Mastermix, 2.5 μL of 10 μM forward primer (BL753), 2.5 μL of 10 μM reverse primer (BL752) in nuclease free water. The reaction was incubated in a thermocycler at 98°C for 3 minutes, then 5 cycles at 98°C for 15 seconds, 67°C for 20 seconds, 72°C for 60 seconds. The pre-amplified products was kept on ice, and 5 μL were used as input for qPCR to determine the optimal number of additional cycles, The qPCR reactions were set up as follows: 5 μL of the pre-amplified material, 5 μL of 2x Q5 master mix, 0.5 μL of 10 μM forward primer (BL753), 0.5 μL of 10 μM reverse primer (BL752), 2.5 μL of 20x EVA-Green in nuclease free water. The cycling protocol was 98°C for 3 minutes, 40 cycles of 98°C for 15 seconds, 67°C for 20 seconds, 72°C for 60 seconds with a plate reading step. The number of cycles required to reach 1/3 of the amplification curve plateau was selected for the final amplification of the libraries. The PCR reactions were purified with AmpureXP beads (0.8x) and eluted in 20 μL of EB buffer. A small aliquot (2 μL) was analysed on a Tapestation D5000 screentape. The amplified cDNA fraction was then tagmented with Tn5 loaded with a single adapter (BL371/BL515) in a reaction set up as follows: 50 ng of pre-amplified cDNA, 0.5 μL loaded Tn5 enzyme in 10 mM Tris-HCl pH 7.6, 5 mM MgCl2, 10% Dimethyl Formamide, 0.33x PBS, 0.1% Tween-20, 0.01% Digitonin in 50uL final. The tagmentation reaction was incubated at 37°C for 30 minutes, and inactivated adding 50 μL of 80 mM EDTA and 2.5 μL of 2% SDS solution for 15 minutes at room temperature. The reactions were purified were purified with AmpureXP beads (0.8x) and eluted in 20 μL of EB buffer. A small aliquot (2 μL) was analysed on a Tapestation. The purified reaction was amplified to generate ATAC libraries as follows: 18 μL of Tn5-cDNA, 2.5 μL of 10uM P5 primer (BL727), 2.5 μL of 10 μM indexed P7 primer (Ad2.X) in nuclease free water. The cycling protocol was 72°C for 5 minutes (gap-fill), then 98°C for 1 minutes and 5 cycles of 98°C for 10 seconds, 68°C for 20 seconds, 72°C for 45 seconds, with a final extension at 72°C for 5 minutes. The PCR reactions were purified with AmpureXP beads (0.8x) and eluted in 20 μL of EB buffer. The libraries were analysed on a Tapestation (D5000 screentape) and quantified with KAPA library quantification kit prior to pooling. Sequencing was performed on a Novaseq (Illumina) instrument.

### BALI barcode demultiplexing

For all sequencing experiments including BALI barcodes (regardless of length), the fastq files produced by high-throughput sequencing were processed through the following pipeline using custom bash shell scripts: 1) UMI extraction and annotation using the umitools package (v0.5.5) with the command *umi_tools extract– -bc-pattern=NNNNNN*, 2) extraction of reads including each first (spatial) BALI barcode from both paired fastq files using the *cutadapt* package (v4.1) (options *-g ^[barcode], -e 1, --pair-filter-both, --match-read-wildcards*), 3) Extraction of successive BALI barcodes from the output of step 2, using the same command and options, 4) Extraction of the final adapter placed before the sequencing library itself (RT oligo for RNAseq, tn5 mosaic end for ATAC-seq) using the same command as above. The strategy was performed iteratively to produce a pair of read1/read2 fastq files for each complete BALI barcode, as well as to produce fastq files for “partial” barcodes caused by incomplete ligations (i.e. in cases in which position 1 of a barcode ligated to the overhang-compatible position 4 rather than to position 2).

Further analysis was performed on all reads featuring full barcodes, as well as on those featuring a complete barcode fragment including position 4 (the spatially defined one) as well as position 3, plus the end adapter (RT or mosaic end). These reads corresponded to molecules that could be reliably assigned to a specific region.

### BALI RNA sequencing analysis

The read 1 fastq file from each spatial region was mapped on the mm10 reference genome (source: Illumina igenomes) using the STAR package (v 2.7.9a) with the following options: *--outReadsUnmapped Fastx – outSAMtype BAM Unsorted–-quantmode GeneCounts*. The resulting BAM files were then sorted and indexed using the *samtools* package (v1.9) commands. To produce UMI counts for each gene (as opposed to the raw counts produced by STAR), each BAM file was run through the *featureCounts* command from the *Subread* package (v2.0.3) with default options against the same mm10 genome annotation described above. The resulting annotated BAM file was sorted again using samtools sort. UMIs counts per gene were calculated using the *umi_tools count* command (from the umitools package v0.5.5) with options–*-per-gene– -gene-tag=XT–-assigned-status-tag=XS,* and UMIs were deduplicated using the *umi_tools group* command (to produce an overall quantification of all UMIs in each region) and *umi_tools dedup* command (to produce a deduplicated BAM file). The deduplicated BAm files were sorted and indexed a third time, and genome coverage was calculated using the *bamCoverage* command from the *deeptools* package (v3.5.1) with the following options–*-normalizeUsing RPKM–-binSize 1*.

For the RNA only experiment on e16.5 mouse embryos, differential expression was performed by merging the UMI count matrices produced by the procedure described above. Each count was normalized by dividing it by the total number of gene-assigned UMIs in the library and multiplying it by 1,000,000, then fold change and log10 change ratios between the SVZ and the cortical areas were calculated using excel. For the multi-omic experiment on adult mouse hippocampus, the UMI count matrices from each area and replicate were merged and normalized as above, and the fold and log10 change ratios calculated using the average expression across replicates.

### Laser capture microdissection data analysis

Raw paired-end sequencing files from each LCM-dissected area were first filtered using the cutadapt package (v4.1) to remove the Illumina sequencing adapters from both the 5’ and 3’ of each sequencing read. The filtered read 1 was then mapped on the mm10 was mapped on the mm10 reference genome (source: Illumina igenomes) using the STAR package (v 2.7.9a) with the following options: *--outReadsUnmapped Fastx –outSAMtype BAM Unsorted --quantmode GeneCounts*. The resulting BAM files were then sorted and indexed using the *samtools* package (v1.9) commands. Genome coverage was calculated using the *bamCoverage* command from the *deeptools* package (v3.5.1) with the following options *--normalizeUsing RPKM --binSize 1.* Differential expression was performed by merging the count matrices produced by the procedure described above. Each count was normalized by dividing it by the total number of gene-assigned reads in the library and multiplying it by 1,000,000, then fold change and log10 change ratios between the SVZ and the cortical areas were calculated using excel.

### Quantification of total UMI per region using existing 10X Visium data for e16.5 embryos

The raw data from Hong, Song, Zhant *et al*.^40^ was obtained from the National Genomics Data Center (https://ngdc.cncb.ac.cn) BioProject accession number PRJCA010498, (sequencing reads under GSA accession number CRA007506, H&E images under OMIX accession number OMIX001298). The raw fastq files and HE image files for replicate 2 of the e16.5 time point were processed using the *Spaceranger* package from 10X genomics (*v2.1.1*) using reference *refdata-gex-mm10-2020-.*The spaceranger output data was processed using the *scanpy* package (v1.9.8) under python. First basic statistics, including percent of reads mapping to mitochondrial genes and total UMIs, were calculated for each spot. Second, spots with <5000 or >35000 UMIs, or with more than 20% Mt reads, were excluded. A custom script (using the *holoviews* and *bokeh* packages) was then used to define, on the high-resolution H&E image associated with the Visium dataset, areas matching the two regions profiled by the BALI RNA experiment. The total number of UMI for each spot within the regions was then calculated and used as a comparison with the BALI data.

### Re-analysis of existing literature data for 10X Visium

The raw data from Vanrobaeys *et al*. was obtained from GEO accession GSE223066. The spaceranger output matrices for samples *Visium-HC5* to *Visium-HC9* (corresponding to “home cage” control conditions) were downloaded and processed using the *scanpy* package (v1.9.8) under python. First basic statistics, including percent of reads mapping to mitochondrial genes and total UMIs, were calculated for each spot. Second, spots with <5000 or >35000 UMIs, or with more than 20% Mt reads, were excluded. A custom script (using the *holoviews* and *bokeh* packages) was then used to define, on the high-resolution H&E image associated with each Visium dataset, areas matching the two regions profiled by the BALI multi-omic experiment. The UMI counts from all the spots included in each of the areas were then summed (pseudobulking), and the resulting count matrices for each replicate dataset (HC5 to HC9) were merged together as a single dataset.

Differential expression was performed as follows: first each count value was normalized by dividing it by the total number of UMIs in that sample/area and multiplying it by 1,000,000, then fold change and log10 change ratios between the DG and CA1 areas were calculated on the average of the UMIs across all replicates for each areas. The fold and log ratios obtained using this method were compared to those obtained by performing normalization and differential expression testing using the R *DeSeq2* package, and showed similar results

### Gene enrichment visualization plots

Gene enrichment were visualized using a custom python script using the following modules: *numpy, holoviews, bokeh*. The script comprised the following sterps: 1) an histological image corresponding to the anatomical areas being measured was used to define two representative polygons corresponding to the profiled areas, 2) the log10 change value for a gene to be visualized was divided by a user-defined value set as maximum color intensity, 3) the resulting value was used to set the color intensity for the polygon in which the gene was upregulated, 4) the two color-graded polygons were drawn overlaid to the anatomical regions of interest. The minimum and maximum scaling values used were −1 and +1 log10 units for LCM data and BALI data on e16.5 embryos, and −0.5 and +0.5 log10 units for multi-omic BALI data and the matching Visium data from *Vanrobaeys et al*.

### ATAC-Seq sequencing analysis

Both read 1 and read 2 files from each spatial region were mapped to the mm10 reference genome (source: Illumina igenomes) using the *bowtie2* package (v2.5.1) using the following options: *--local --very-sensitive-local --soft-clipped-unmapped-tlen --dovetail --no-mixed -q --no-discordant --phred33 -I 10 -X 1000*. The resulting SAM files were converted to BAM, sorted and indexed using the *samtools* package (v1.9). Genome coverage was calculated using the *bamCoverage* command from the *deeptools* package (v3.5.1) with the following options *-normalizeUsing None --binSize 3 --scaleFactor [scale_factor]*. The scale factor was calculated as *1 / total mapped reads* for each library.

Peaks were identified using the macs2 package (v2.2.9.1) using the following options *-f BAMPE –nomodel --keep-dup=all*.

### Re-analysis of existing literature data for integrated scRNAseq/scATACseq

Vanrobaeys *et al.*^47^ includes matched single-cell RNA-seq (produced using the split-seq protocol from Parse Biosciences) and single-cell ATAC-seq (produced using the 10X ATAC-seq protocol) for the hippocampus of control mice (home cage condition). We used this data to identify cellular populations corresponding to the dentate gyrus granule cells and CA1 pyramidal cells, and match their genome coverage profile to the one obtained by BALI in the corresponding areas.

All analysis was performed in R using packages *AnnotationHub (v3.6.0), Seurat (v5.0.1) and Signac (v1.12.9004)*. Annotation data was extracted from the EnsemblDB Mus Musculus annotation v98 (corresponding to the annotation used by the cellranger-ATAC reference genome used in the paper). scRNAseq analysis and scATACseq analysis were performed as described in the “Split-Seq Analysis” and “Single Nuclei ATAC-Sequencing data analysis” sections of Vanrobaeys *et al*, including integration between the two datasets. The outputs from the cellranger-ATAC pipeline and split-pipe pipeline were downloaded from GEO accession GSE223066. For ATAC-Seq, the two replicate datasets were merged before dimensionality reduction and clustering.

The CA1 and DG populations were detected in the scRNAseq data by observing overexpression of several markers including *C1ql2, Neurod2, Synpr, Prox1* (DG) and *Mpped1* and *Neurod6* (CA1 pyramidal). Genome coverage plots were obtained for the clusters in the scATACSeq data for which the predicted transfer label from scRNAseq matched the CA1 and DG populations using the Signac *CoveragePlot* function.

