## Supplementary material for "Spatially tuneable multi-omics sequencing using light-driven combinatorial barcoding of molecules in tissues": Consortium Author List

| Surname | First name | Role | ORCid | Affiliation |
|---|---|---|---|---|
| **Hannon** | **Gregory J** | **Co-I** | **https://orcid.org/0000-0003-4021-3898** | **Cancer Research UK Cambridge Institute, Li Ka Shing Centre, University of Cambridge, Cambridge CB2 0RE, UK** |
| Albuquerque | Bruno | Postdoc | https://orcid.org/0000-0003-1257-9120 | Cancer Research UK Cambridge Institute, Li Ka Shing Centre, University of Cambridge, Cambridge CB2 0RE, UK |
| Alini | Martina | Research Assistant | | Cancer Research UK Cambridge Institute, Li Ka Shing Centre, University of Cambridge, Cambridge CB2 0RE, UK |
| Ashmore | Heather | Project Manager | | Cancer Research UK Cambridge Institute, Li Ka Shing Centre, University of Cambridge, Cambridge CB2 0RE, UK |
| Ashmore | Thomas | Research Assistant | | Cancer Research UK Cambridge Institute, Li Ka Shing Centre, University of Cambridge, Cambridge CB2 0RE, UK |
| Battistoni | Giorgia | Postdoc | https://orcid.org/0000-0003-1257-9120 | Cancer Research UK Cambridge Institute, Li Ka Shing Centre, University of Cambridge, Cambridge CB2 0RE, UK |
| **Bressan** | **Dario** | **Co-I, IMAXT Lab Head** | **https://orcid.org/0000-0003-3592-699X** | **Cancer Research UK Cambridge Institute, Li Ka Shing Centre, University of Cambridge, Cambridge CB2 0RE, UK** |
| Cannell | Ian Gordon | Senior Research Associate | https://orcid.org/0000-0001-5832-9210 | Cancer Research UK Cambridge Institute, Li Ka Shing Centre, University of Cambridge, Cambridge CB2 0RE, UK |
| Casbolt | Hannah | Research Assistant | | Cancer Research UK Cambridge Institute, Li Ka Shing Centre, University of Cambridge, Cambridge CB2 0RE, UK |
| Deighton | Lauren | PhD student | | Cancer Research UK Cambridge Institute, Li Ka Shing Centre, University of Cambridge, Cambridge CB2 0RE, UK |
| Falciatori | Ilaria | Project Manager | | |
| Boquetale | Carla | Research Assistant | | Cancer Research UK Cambridge Institute, Li Ka Shing Centre, University of Cambridge, Cambridge CB2 0RE, UK |
| Coutts | Nikki | Project Manager | | |
| Ying Sia | Chee | Research Assistant | | |
| Fatemi | Atefeh | Research Assistant | | Cancer Research UK Cambridge Institute, Li Ka Shing Centre, University of Cambridge, Cambridge CB2 0RE, UK |
| Hemmer | Nicole | Research Assistant | | Cancer Research UK Cambridge Institute, Li Ka Shing Centre, University of Cambridge, Cambridge CB2 0RE, UK |
| Hua | Kui | Postdoc | | |
| Jauset | Cristina | PhD student | https://orcid.org/0000-0003-1408-026X | Cancer Research UK Cambridge Institute, Li Ka Shing Centre, University of Cambridge, Cambridge CB2 0RE, UK |
| Kovačević | Tatjana | PhD student | | Cancer Research UK Cambridge Institute, Li Ka Shing Centre, University of Cambridge, Cambridge CB2 0RE, UK |
| Mulvey | Claire M | Principal Research Associate | https://orcid.org/0000-0002-2989-2052 | Cancer Research UK Cambridge Institute, Li Ka Shing Centre, University of Cambridge, Cambridge CB2 0RE, UK |
| Narayanan | Natasha | Mphil Student | | |
| Nugent | Fiona | PhD student | https://orcid.org/0000-0001-8148-0867 | Cancer Research UK Cambridge Institute, Li Ka Shing Centre, University of Cambridge, Cambridge CB2 0RE, UK |
| Rebbeck | Clare | Senior Research Associate | | |
| Paez Ribes | Marta | Principal Research Associate | | Cancer Research UK Cambridge Institute, Li Ka Shing Centre, University of Cambridge, Cambridge CB2 0RE, UK |
| Pearsall | Isabella | PhD student | https://orcid.org/0000-0001-7771-3082 | Cancer Research UK Cambridge Institute, Li Ka Shing Centre, University of Cambridge, Cambridge CB2 0RE, UK |
| Pearsall | Sarah | Postdoc | | |
| Qosaj | Fatime | PhD student | | Cancer Research UK Cambridge Institute, Li Ka Shing Centre, University of Cambridge, Cambridge CB2 0RE, UK |
| Sawicka | Kirsty | Postdoc | https://orcid.org/0000-0003-4195-6327 | Cancer Research UK Cambridge Institute, Li Ka Shing Centre, University of Cambridge, Cambridge CB2 0RE, UK |
| Wild | Sophia A | PhD student | https://orcid.org/0000-0003-0397-6255 | Cancer Research UK Cambridge Institute, Li Ka Shing Centre, University of Cambridge, Cambridge CB2 0RE, UK |
| Williams | Elena | Research Assistant | | |
| **Ali** | **Hamid Raza** | **Co-I** | **https://orcid.org/0000-0001-7587-0906** | **Cancer Research UK Cambridge Institute, Li Ka Shing Centre, University of Cambridge, Cambridge CB2 0RE, UK** |
| **Aparicio** | **Samuel** | **Co-I** | **https://orcid.org/0000-0002-0487-9599** | **Department of Molecular Oncology, BC Cancer, part of the Provincial Health Services Authority, Vancouver, BC, Canada; Department of Pathology and Laboratory Medicine, University of British Columbia, Vancouver, BC, Canada** |
| Laks | Emma | PhD student | | Department of Molecular Oncology, BC Cancer, part of the Provincial Health Services Authority, Vancouver, BC, Canada; Department of Pathology and Laboratory Medicine, University of British Columbia, Vancouver, BC, Canada |
| Li | Yangguang | Postdoc | | Department of Molecular Oncology, BC Cancer, part of the Provincial Health Services Authority, Vancouver, BC, Canada |
| O'Flanagan | Ciara H | Research Associate | https://orcid.org/0000-0002-4570-7347 | Department of Molecular Oncology, BC Cancer, part of the Provincial Health Services Authority, Vancouver, BC, Canada |
| Smith | Austin | Undergraduate student | | Department of Molecular Oncology, BC Cancer, part of the Provincial Health Services Authority, Vancouver, BC, Canada |
| Ruiz | Teresa | Technician | | Department of Molecular Oncology, BC Cancer, part of the Provincial Health Services Authority, Vancouver, BC, Canada |
| Lai | Daniel | Senior Bioinformatics Scientist | https://orcid.org/0000-0001-9203-6323 | Department of Molecular Oncology, BC Cancer, part of the Provincial Health Services Authority, Vancouver, BC, Canada; Department of Pathology and Laboratory Medicine, University of British Columbia, Vancouver, BC, Canada |
| Roth | Andrew | IMAXT Affiliate | https://orcid.org/0000-0003-3422-8823 | Department of Molecular Oncology, BC Cancer, part of the Provincial Health Services Authority and Department of Computer Science and Department of Pathology and Laboratory Medicine, University of British Columbia, Vancouver, BC, Canada. |
| Au | Vinci | Research Assistant | https://orcid.org/0000-0002-3587-6200 | Department of Molecular Oncology, BC Cancer, part of the Provincial Health Services Authority, Vancouver, BC, Canada |
| Baril | Caroline | Research Assistant | | Department of Molecular Oncology, BC Cancer, part of the Provincial Health Services Authority, Vancouver, BC, Canada |
| Beatty | Sean | Bioinformatics Scientist | https://orcid.org/0000-0001-6819-7071 | Department of Molecular Oncology, BC Cancer, part of the Provincial Health Services Authority, Vancouver, BC, Canada |
| **Balasubramanian** | **Shankar** | **Co-I** | | **Cancer Research UK Cambridge Institute, Li Ka Shing Centre, University of Cambridge, Cambridge CB2 0RE, UK; Department of Chemistry, University of Cambridge, Lensfield Road, Cambridge, CB2 1EW, UK; School of Clinical Medicine, University of Cambridge, Cambridge, CB2 0SP, UK.** |
| Nogueira | João CF | Postdoc | | Cancer Research UK Cambridge Institute, Li Ka Shing Centre, University of Cambridge, Cambridge CB2 0RE, UK; Department of Chemistry, University of Cambridge, Lensfield Road, Cambridge, CB2 1EW, UK |
| Lee | Max | Postdoc | | |
| **Bodenmiller** | **Bernd** | **Co-I** | | **Institute of Molecular Life Sciences, University of Zurich, Zurich 8057, Switzerland --> as of 01-01-20 change to Department of Quantitative Biomedicine, , University of Zurich, Zurich 8057, Switzerland** |
| Bollhagen | Alina | PhD student | | Department of Quantitative Biomedicine, University of Zurich, Zurich 8057, Switzerland |
| Burger | Marcel | Postdoc | https://orcid.org/0000-0002-9904-2547 | Institute of Molecular Life Sciences, University of Zurich, Zurich 8057, Switzerland --> as of 01-01-20 change to Department of Quantitative Biomedicine, , University of Zurich, Zurich 8057, Switzerland |
| Kuett | Laura | PhD student | | Institute of Molecular Life Sciences, University of Zurich, Zurich 8057, Switzerland --> as of 01-01-20 change to Department of Quantitative Biomedicine, , University of Zurich, Zurich 8057, Switzerland |
| Windhager | Jonas | PhD student | https://orcid.org/0000-0002-2111-5291 | Institute of Molecular Life Sciences, University of Zurich, Zurich 8057, Switzerland --> as of 01-01-20 change to Department of Quantitative Biomedicine, , University of Zurich, Zurich 8057, Switzerland |
| **Boyden** | **Edward S** | **Co-I** | | **McGovern Institute, Departments of Biological Engineering and Brainand Cognitive Sciences, Massachusetts Institute of Technology,Cambridge, Massachusetts, USA, and HHMI, Cambridge, Massachusetts, USA** |
| Ghosh | Debarati | Research Staff | | McGovern Institute, Departments of Biological Engineering and Brainand Cognitive Sciences, Massachusetts Institute of Technology,Cambridge, Massachusetts, USA, and HHMI, Cambridge, Massachusetts, USA |
| Sinha | Anubhav | PhD student | | McGovern Institute, Departments of Biological Engineering and Brainand Cognitive Sciences, Massachusetts Institute of Technology,Cambridge, Massachusetts, USA, and HHMI, Cambridge, Massachusetts, USA |
| Pryor | Brett | Research Assistant | | McGovern Institute, Departments of Biological Engineering and Brainand Cognitive Sciences, Massachusetts Institute of Technology,Cambridge, Massachusetts, USA, and HHMI, Cambridge, Massachusetts, USA |
| Zhang | Ruihan | PhD student | | McGovern Institute, Departments of Biological Engineering and Brainand Cognitive Sciences, Massachusetts Institute of Technology,Cambridge, Massachusetts, USA, and HHMI, Cambridge, Massachusetts, USA |

| | | | | |
|---|---|---|---|---|
| Lovell | Jack | Research Assistant | | McGovern Institute, Departments of Biological Engineering and Brainand Cognitive Sciences, Massachusetts Institute of Technology,Cambridge, Massachusetts, USA, and HHMI, Cambridge, Massachusetts, USA |
| Zhang | Chi | Research Staff | | McGovern Institute, Departments of Biological Engineering and Brainand Cognitive Sciences, Massachusetts Institute of Technology,Cambridge, Massachusetts, USA, and HHMI, Cambridge, Massachusetts, USA |
| Lu | Yangning | Postdoc | | McGovern Institute, Departments of Biological Engineering and Brainand Cognitive Sciences, Massachusetts Institute of Technology,Cambridge, Massachusetts, USA, and HHMI, Cambridge, Massachusetts, USA |
| **Caldas** | **Carlos** | **Co-I** | | **Department of Oncology and Cancer Research UK Cambridge Institute, University of Cambridge, Cambridge, CB2 0RE, UK** |
| Bruna | Alejandra | Postdoc | | Department of Oncology and Cancer Research UK Cambridge Institute, University of Cambridge, Cambridge, CB2 0RE, UK |
| Callari | Maurizio | Postdoc | | Cancer Research UK Cambridge Institute, Li Ka Shing Centre, University of Cambridge, Cambridge CB2 0RE, UK |
| Deighton | Lauren | PhD student | | Cancer Research UK Cambridge Institute, Li Ka Shing Centre, University of Cambridge, Cambridge CB2 0RE, UK |
| Greenwood | Wendy | RA | | Cancer Research UK Cambridge Institute, Li Ka Shing Centre, University of Cambridge, Cambridge CB2 0RE, UK |
| Lerda | Giulia | PhD student | | Cancer Research UK Cambridge Institute, Li Ka Shing Centre, University of Cambridge, Cambridge CB2 0RE, UK |
| Eyal-Lubling | Yaniv | Postdoc | | Department of Oncology and Cancer Research UK Cambridge Institute, University of Cambridge, Cambridge, CB2 0RE, UK |
| Rueda | Oscar M | Postdoc | | Department of Oncology and Cancer Research UK Cambridge Institute, University of Cambridge, Cambridge, CB2 0RE, UK |
| Shea | Abigail | PhD Student | | Department of Oncology and Cancer Research UK Cambridge Institute, University of Cambridge, Cambridge, CB2 0RE, UK |
| **Harris** | **Owen** | **Co-I** | | **Súil Interactive Ltd, Dame Lane, Dublin, UK** |
| Becker | Robby | Lead Programmer | | Súil Interactive Ltd, Dame Lane, Dublin, UK |
| Duncan | Natalie | Project Administrator | | Súil Interactive Ltd, Dame Lane, Dublin, UK |
| Grimaldi | Flaminia | Lead Artist | | Súil Interactive Ltd, Dame Lane, Dublin, UK |
| Harris | Suvi | Liaison with science groups | | Súil Interactive Ltd, Dame Lane, Dublin, UK |
| Vogl | Sara Lisa | VR Designer | | Súil Interactive Ltd, Dame Lane, Dublin, UK |
| Weselak | Joanna | Animation Specialist | | Súil Interactive Ltd, Dame Lane, Dublin, UK |
| **Joyce** | **Johanna A** | **Co-I** | **https://orcid.org/0000-0002-6332-2598** | **Department of Oncology and Ludwig Institute for Cancer Research, University of Lausanne, Lausanne, Switzerland** |
| Watson | Spencer S | Postdoc | https://orcid.org/0000-0002-5583-1544 | Department of Oncology and Ludwig Institute for Cancer Research, University of Lausanne, Lausanne, Switzerland |
| **Marioni** | **John** | **Co-I** | **https://orcid.org/0000-0001-9092-0852** | **Cancer Research UK Cambridge Institute, Li Ka Shing Centre, University of Cambridge, Cambridge CB2 0RE, UK; EMBL-European Bioinformatics Institute, Wellcome Genome Campus, Cambridge, UK; Wellcome Sanger Institute, Wellcome Genome Campus, Cambridge, UK** |
| **Shah** | **Sohrab P** | **Co-I** | **https://orcid.org/0000-0001-6402-523X** | **Department of Molecular Oncology, British Columbia Cancer Research Centre, Vancouver, BC, Canada; Department of Pathology and Laboratory Medicine, University of British Columbia, Vancouver, BC, Canada; Computational Oncology, Department of Epidemiology and Biostatistics, Memorial Sloan Kettering Cancer Center, New York, USA** |
| McPherson | Andrew | Bioinformatician | | Department of Molecular Oncology, British Columbia Cancer Research Centre, Vancouver, BC, Canada; Computational Oncology, Department of Epidemiology and Biostatistics, Memorial Sloan Kettering Cancer Center, New York, USA |
| Vázquez-García | Ignacio | Postdoc | https://orcid.org/0000-0003-0427-2639 | Computational Oncology, Department of Epidemiology and Biostatistics, Memorial Sloan Kettering Cancer Center, New York, USA; Herbert and Florence Irving Institute for Cancer Dynamics, Columbia University, New York, NY, USA |
| **Tavaré** | **Simon** | **Co-I** | **https://orcid.org/0000-0002-3716-4952** | **Cancer Research UK Cambridge Institute, Li Ka Shing Centre, University of Cambridge, Cambridge CB2 0RE, UK; Herbert and Florence Irving Institute for Cancer Dynamics, Columbia University, New York, NY, USA; New York Genome Center, New York, NY, USA** |
| Dinh | Khanh N | Associate Research Scientist | | Herbert and Florence Irving Institute for Cancer Dynamics, Columbia University, New York, NY, USA |
| Kunes | Russell | PhD student | | Herbert and Florence Irving Institute for Cancer Dynamics, Columbia University, New York, NY, USA |
| **Walton** | **Nicholas A** | **Co-I** | **https://orcid.org/0000-0003-3983-8778** | **Institute of Astronomy, University of Cambridge, Madingley Road, Cambridge, CB3 0HA, UK** |
| Al Sa'd | Mohammad | Postdoc | | Institute of Astronomy, University of Cambridge, Madingley Road, Cambridge, CB3 0HA, UK |
| Chornay | Nick | PhD student | | Institute of Astronomy, University of Cambridge, Madingley Road, Cambridge, CB3 0HA, UK |
| Dariush | Ali | Postdoc | | Institute of Astronomy, University of Cambridge, Madingley Road, Cambridge, CB3 0HA, UK |
| González-Solares | Eduardo A | Postdoc | | Institute of Astronomy, University of Cambridge, Madingley Road, Cambridge, CB3 0HA, UK |
| González-Fernández | Carlos | Postdoc | | Institute of Astronomy, University of Cambridge, Madingley Road, Cambridge, CB3 0HA, UK |
| Irfan | Melis | Postdoc | | Institute of Astronomy, University of Cambridge, Madingley Road, Cambridge, CB3 0HA, UK |
| Küpcü Yoldaş | Aybüke | Postdoc | | Institute of Astronomy, University of Cambridge, Madingley Road, Cambridge, CB3 0HA, UK |
| Molaeinezhad | Alireza | Postdoc | | Institute of Astronomy, University of Cambridge, Madingley Road, Cambridge, CB3 0HA, UK |
| Millar | Neil | Technician | | Institute of Astronomy, University of Cambridge, Madingley Road, Cambridge, CB3 0HA, UK |
| Smith | Leigh | Postdoc | | Institute of Astronomy, University of Cambridge, Madingley Road, Cambridge, CB3 0HA, UK |
| Whitmarsh | Tristan | Postdoc | | Institute of Astronomy, University of Cambridge, Madingley Road, Cambridge, CB3 0HA, UK |
| **Zhuang** | **Xiaowei** | **Co-I** | | **Howard Hughes Medical Institute, Harvard University, Cambridge, MA 02138, USA; Department of Physics, Harvard University, Cambridge, MA 02138, USA; Department of Chemistry and Chemical Biology, Harvard University, Cambridge, MA 02138, USA** |
| Fan | Jean | Postdoc | | Howard Hughes Medical Institute, Harvard University, Cambridge, MA 02138, USA; Department of Physics, Harvard University, Cambridge, MA 02138, USA; Department of Chemistry and Chemical Biology, Harvard University, Cambridge, MA 02138, USA |
| Lee | Hsuan | PhD student | | Howard Hughes Medical Institute, Harvard University, Cambridge, MA 02138, USA; Department of Physics, Harvard University, Cambridge, MA 02138, USA; Department of Chemistry and Chemical Biology, Harvard University, Cambridge, MA 02138, USA |
| Sepúlveda | Leonardo A | Postdoc | https://orcid.org/0000-0003-3602-1009 | Howard Hughes Medical Institute, Harvard University, Cambridge, MA 02138, USA; Department of Physics, Harvard University, Cambridge, MA 02138, USA; Department of Chemistry and Chemical Biology, Harvard University, Cambridge, MA 02138, USA |
| Xia | Chenglong | PhD student | https://orcid.org/0000-0002-5895-6342 | Howard Hughes Medical Institute, Harvard University, Cambridge, MA 02138, USA; Department of Physics, Harvard University, Cambridge, MA 02138, USA; Department of Chemistry and Chemical Biology, Harvard University, Cambridge, MA 02138, USA |
| Zheng | Pu | PhD student | | Howard Hughes Medical Institute, Harvard University, Cambridge, MA 02138, USA; Department of Physics, Harvard University, Cambridge, MA 02138, USA; Department of Chemistry and Chemical Biology, Harvard University, Cambridge, MA 02138, USA |
