## Supplementary Figures for "Spatially tuneable multi-omics sequencing using light-driven combinatorial barcoding of molecules in tissues"

SupplementaryFigure 1

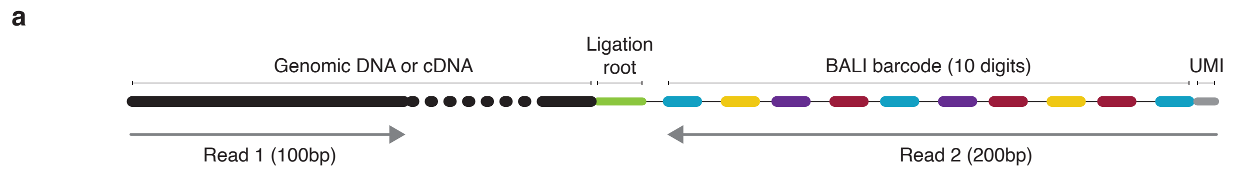


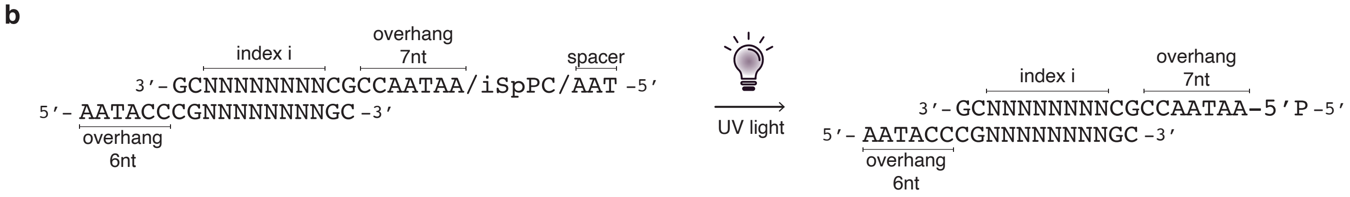

**Supplementary Figure 1. BALI library and index structure**

a) *Library structure*. Genomic or 3'end terminal of the cDNA are shown in black (dotted region to represent variable fragment sizes). The ligation root is represented in green. The BALI spatial barcode is exemplified as a 10-digit barcode, where each digit is shown as different colour box. A region with the Unique Molecular Identifier is shown as a grey box. Paired end sequencing is used to read the cDNA/genomic fragment sequence (Read 1, 100 bp) and the spatial barcode plus UMI (Read 2, 200 bp). b) *Index structure.* Each index cassette is composed of two staggered complementary oligonucleotides. On the extending strand (upper) there is a variable sequence that corresponds to each individual index value, flanked by GC staples to ensure strong annealing at the ligation site. At the 5'end there is a ligation overhang (alternating between a fixed 7nt or 6nt sequence), a photo-cleavable spacer, and a terminal 5' spacer (5'-AAT-3'). On the opposite strand, there is the reverse complemented sequence to the index and staples, plus the reverse complement sequence for the orthogonal overhang (6-nt if 7nt on the leading strand, or the opposite). Upon illumination with UV light, the photo-cleavable spacer is released to leave a 5' phosphate available for T4 DNA ligase mediated ligation.

Supplementary Figure 2

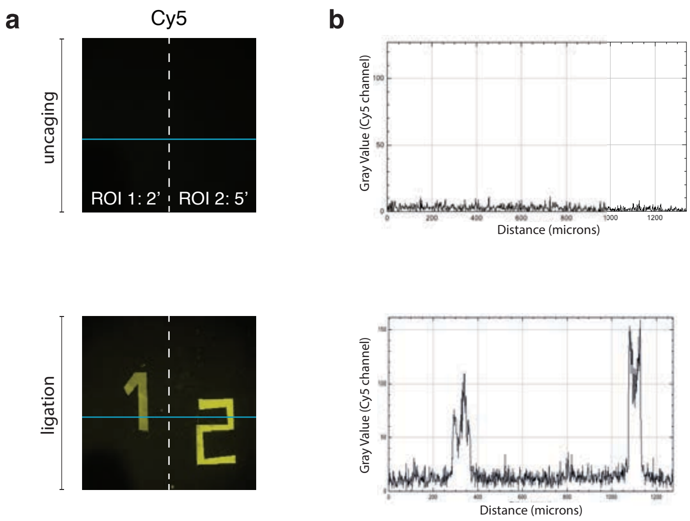

**Supplementary Figure 2. Overhang screen and validation**

a) *Experimental set-up.* A library of oligos with variable 5' ligation overhangs and terminal Illumina P5 and P7 sequences were annealed to produce double-strand DNA indices and ligated as illustrated in figure. The resulting molecules were directly sequenced through Illumina sequencing. Overhangs with different lengths, 6 or 7 nucleotides, where processed separately. b) *Bar plots representing the frequency of reads for each overhang over the total number of reads.* Overhangs were ranked by frequency (highest to lowest from left to right). The colour of the bars represents the GC content of the overhang. Average across 3 replicates shown. Error bars not shown for ease of representation. c, d). *In situ validation of the efficiency validation for the best and worst overhangs identified in the screen for 6-mers (c) and 7mers (d).* The efficiency ligation was measured by densitometry on a denaturing Urea PAGE gel. The individual efficiencies are shown as bar plots, the average efficiencies are shown as box plots (central line for average, whiskers to show min and max values, box extends from $25^{th}$ to $75^{th}$ percentile). Blue for 'best' candidates, red for 'worst' candidates. The chosen overhangs are in darker blue and labelled with an arrow in the bar plot.

Supplementary Figure 3

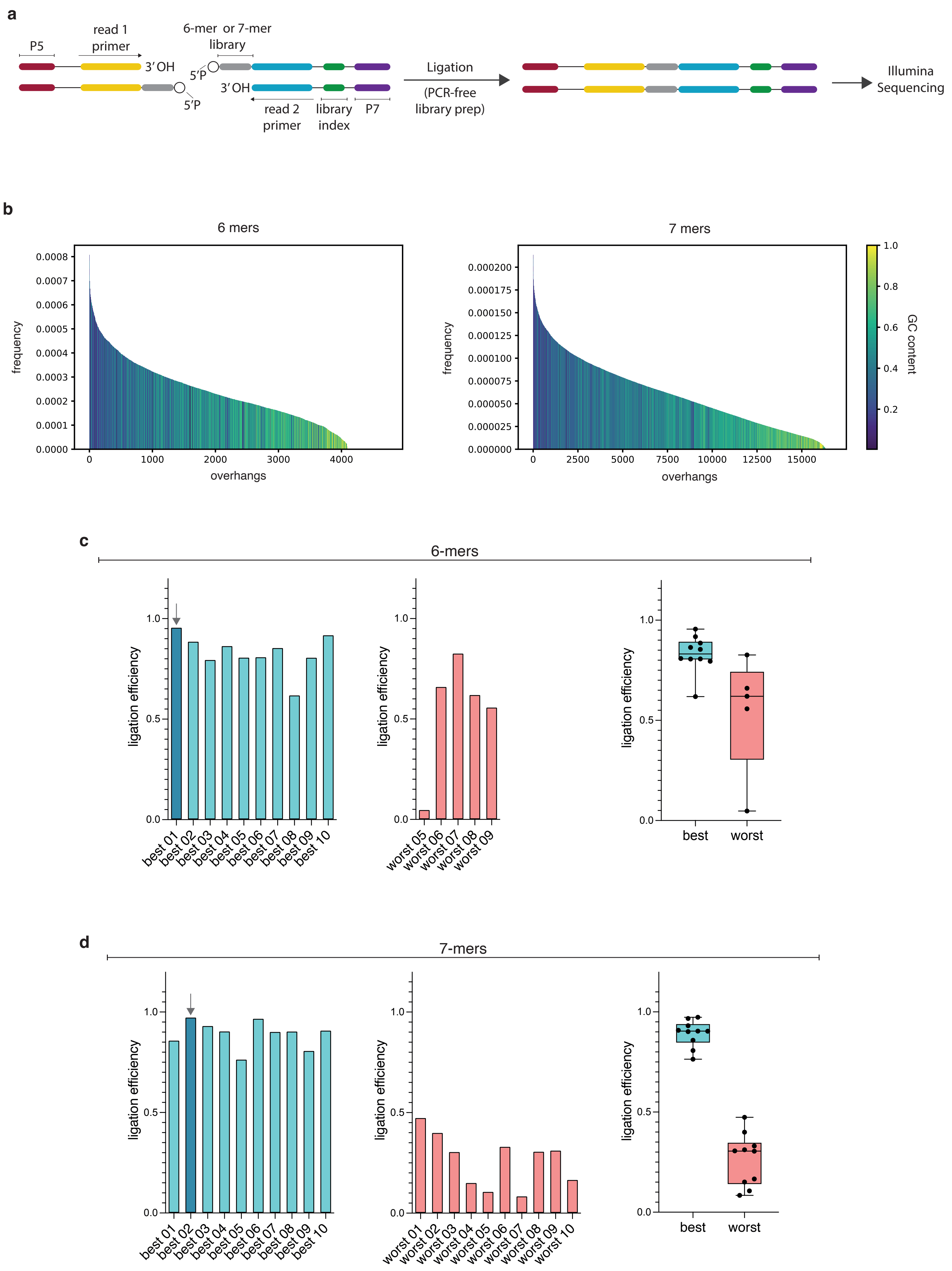

**Supplementary Figure 3. Traces for spatial ligation on solid surface**

a) Confocal microscopy images to show the fluorescence signal derived from the ligated index (Cy5 labelled), after uncaging and ligation. b) Fluorescence signal profiles across a given pixel row (in blue) to illustrate specific ligation and high signal-to-noise.

Supplementary Figure 4

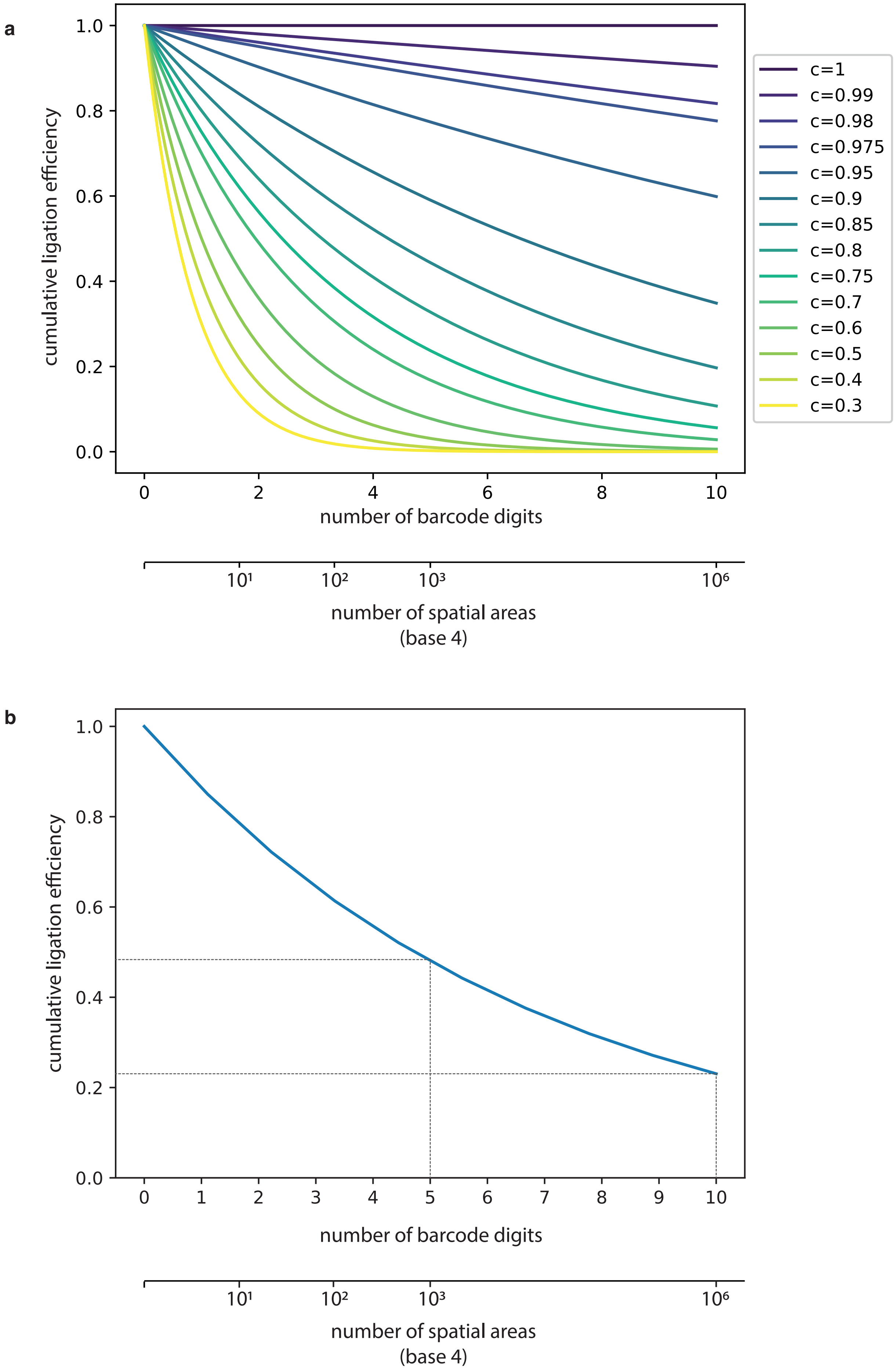

**Supplementary Figure 4. Projections for the cumulative efficiency of ligation for multi-digit barcodes.** a) Projection of the cumulative efficiency of ligation after *n* sequential cycles depending on the average efficiency per cycle (c value = 0.3÷1). The double x-axis represents the number of cycles (*n*), and the corresponding maximum number of spatial barcodes with *n* digits and 4 different values available at each digit (base 4). b) Projection of the cumulative efficiency of after n sequential cycles based on the average efficiency per cycle measured in tissue, c=0.8635 (Fig. 2c). The double x-axis represents the number of cycles (*n*), and the corresponding maximum number of spatial barcodes with *n* digits and 4 different values available at each digit (base 4). The intercepts represent the projected cumulative efficiencies requires to encode ~ 1000 spatial areas (5 digits, base 4) or ~1 million (10 digits, base 4).

Supplementary Figure 5

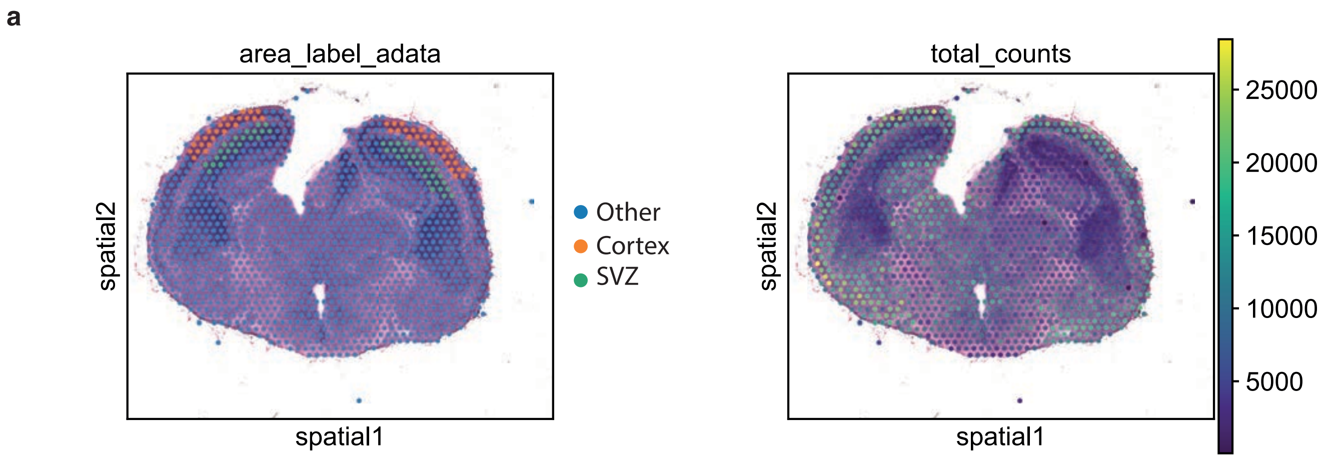

**Supplementary Figure 5. Re-analysis of comparable 10x Visium dataset from E16.5 mouse brain.** H&E image overlayed with the location of the 10x Visium barcode areas[40]. Spots are color-coded to illustrate a) the areas that have been pseudo-bulked as "SVZ" and "cortex" in our analysis or b) UMIs/spot.

Supplementary Figure 6

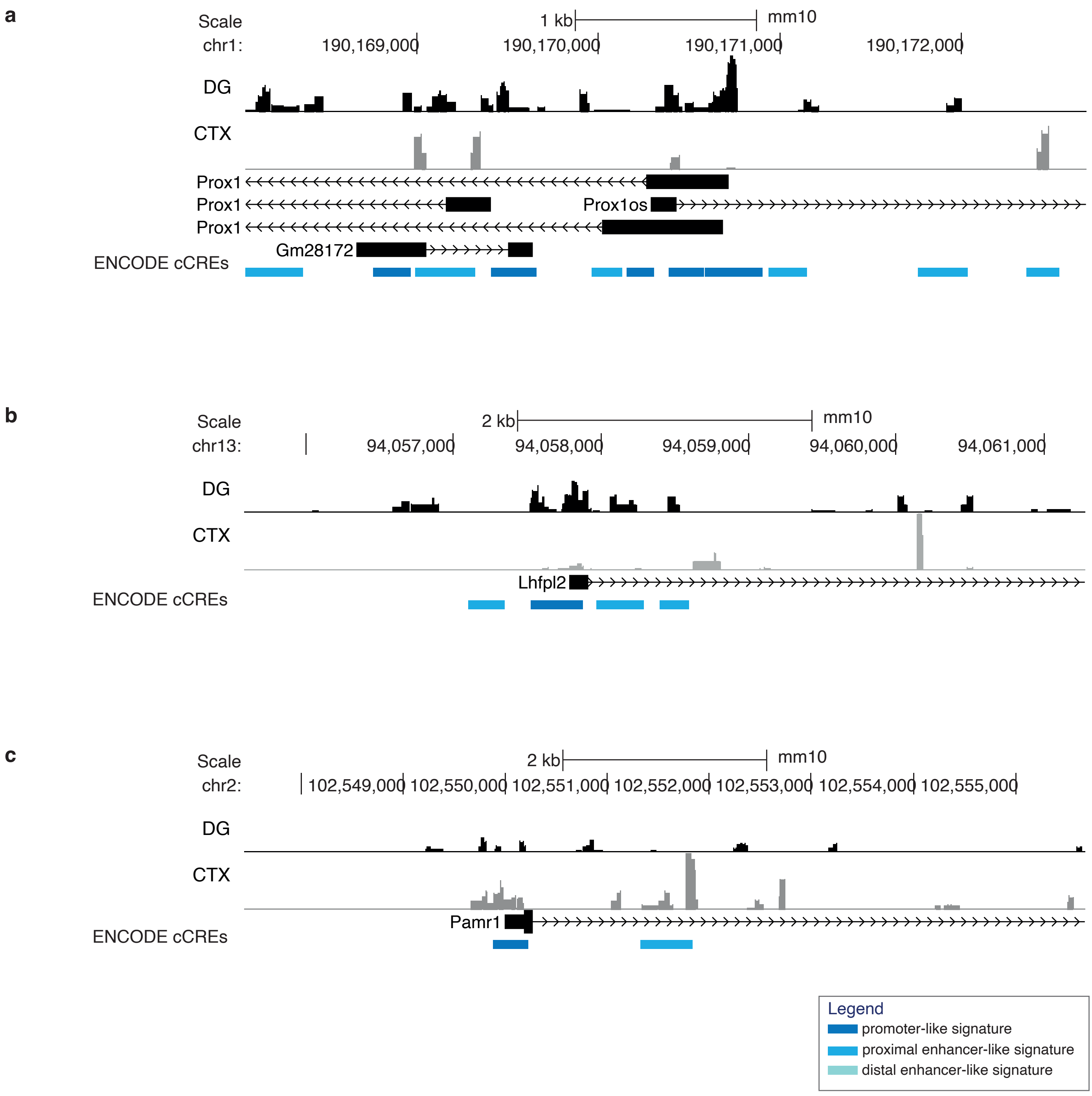

**Supplementary Figure 6. Coverage plots for chromatin accessibility at additional TSSs**

Accessibility for the DG and CTX are shown separately in black and grey, respectively. The RefSeq annotation is shown below, as well the annotation for regulatory elements from the ENCODE's cCREs database[46] a) Prox1, b) Lhfpl2, c) Pamr1

Supplementary Figure 7

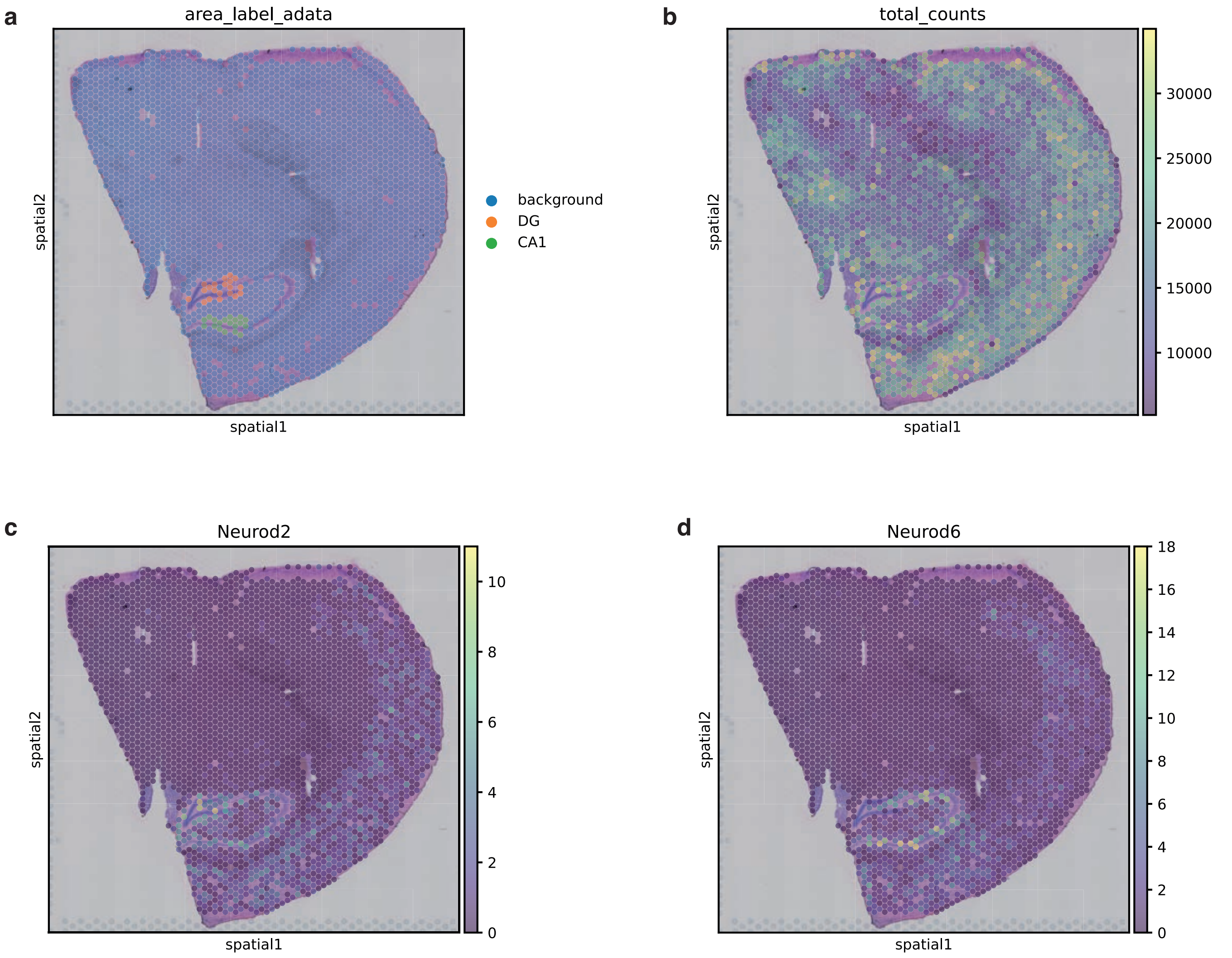

**Supplementary Figure 7. Re-analysis of comparable 10x Visium dataset from adult brain (hippocampus).**

H&E image overlayed with the location of the 10x Visium barcode areas[47]. Spots are color-coded to illustrate a) the areas that have been pseudo-bulked as DG and CA1 in our analysis, b) UMIs/spot, c) expression levels for Neurod2 and d) expression levels for Neurod6

Supplementary Figure 8

**a**

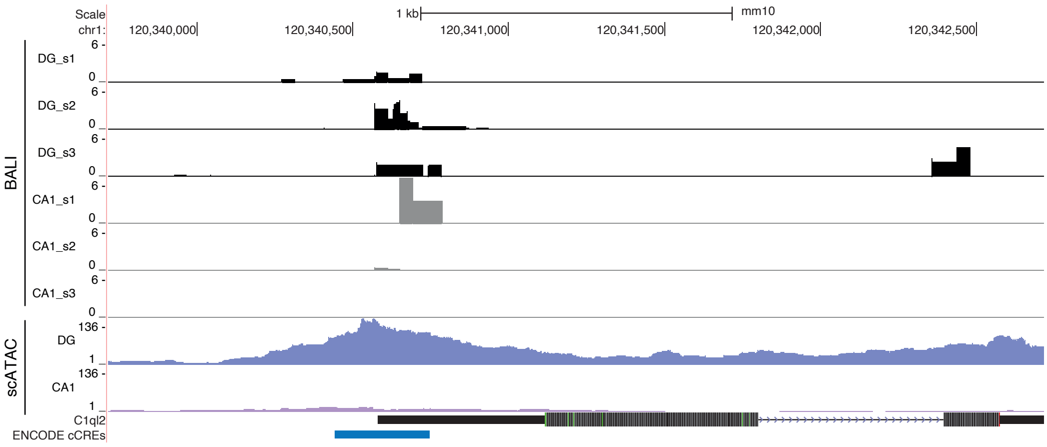


**b**

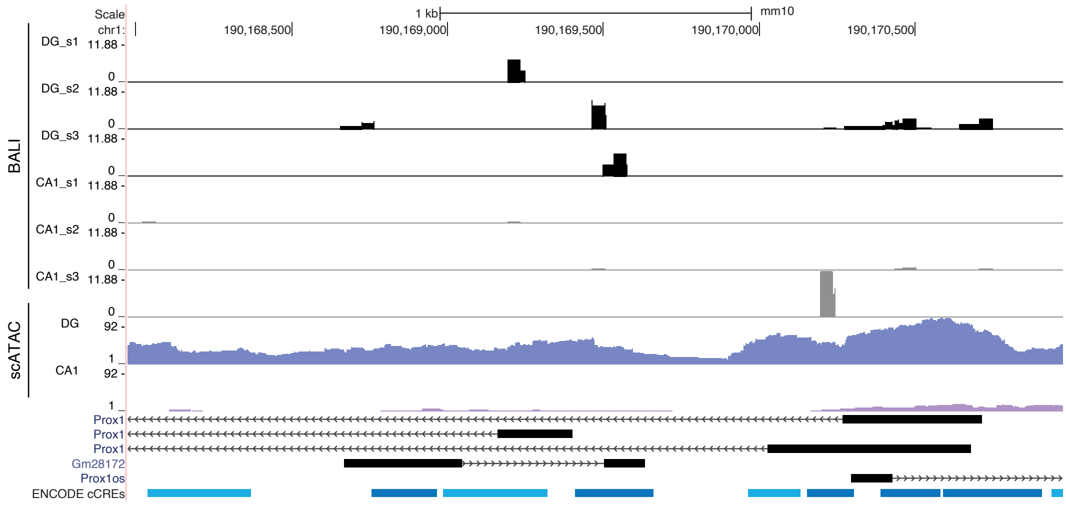


**c**

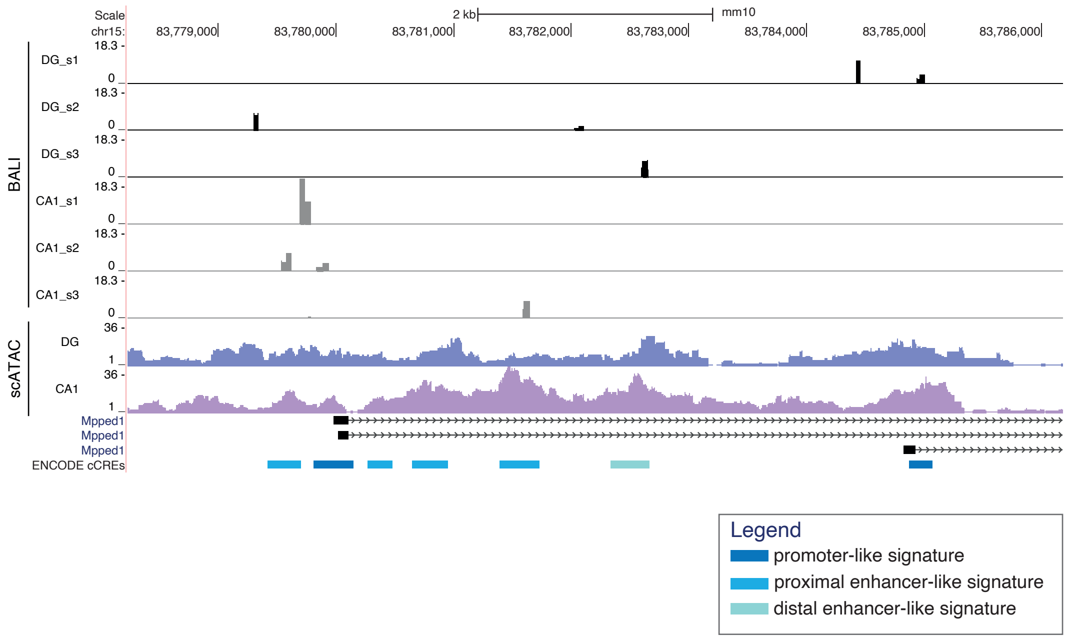

**Supplementary Figure 8. Coverage plots for multiomics profiling at additional TSSs.**
Accessibility for the DG and CA1 are shown for all replicates separately, and for pseudo-bulked disaggregated scATAC from a comparable sample[47]. For BALI, the DG and CA1 are shown in black and grey, respectively. For sc-ATAC, the DG and CA1 are shown in purple and pink, respectively. The RefSeq annotation is shown below, as well the annotation for regulatory elements from the ENCODE's cCREs database[47] a) C1ql2, b) Prox1, c) Mpped1
